# A molecular description of plant cellulose biosynthesis inhibition

**DOI:** 10.64898/2026.07.20.739232

**Authors:** Louis F. L. Wilson, Chaemyeong Lim, Miguel Ángel Torres, Steve Scheiner, Yueping Wan, Pallinti Purushotham, Ruoya Ho, Jochen Zimmer

## Abstract

Dictating cell growth and morphology, cellulose biosynthesis is intrinsic to plant cell biology. Accordingly, cellulose biosynthesis inhibitors (CBIs) are important herbicides, toxins, and experimental tools. We currently lack mechanistic understanding of CBI activity, preventing engineering of herbicide selectivity and disease immunity. Contrasting classical inhibitors, we unexpectedly identify the unusual *Streptomyces* phytotoxin thaxtomin A as the only *in vitro*-active CBI, with unprecedentedly broad-spectrum activity against various cellulose synthase enzymes. High-resolution cryo-electron microscopy reveals that thaxtomin A leverages exotic nitroaromatic chemistry to target a strictly conserved site in cellulose synthase’s polysaccharide secretion channel. Strikingly, *in vitro* biosynthesis and biophysical assays demonstrate thaxtomin A’s near-picomolar efficacy. Plant and algal systems reveal that its global arrest of cellulose biosynthesis produces an osmotically driven crisis in expanding cells. Finally, site-directed mutagenesis generates the first toxin-resistant cellulose synthase. Our results underscore cellulose’s critical function in plant lifeforms and inform efforts to inhibit related enzymes across kingdoms.

## Introduction

Plant cells exist under osmotically generated hydrostatic pressure. Cell walls perform the essential role of controlling and directing this pressure to dictate growth and cell form^1,2^. This tenet is thought to guide all aspects of plant development and viability.

The load-bearing strength of all land plant cell walls is likely provided by cellulose microfibrils^2^. With their orientation leading the direction of cell expansion, microfibrils are directionally deposited in a poorly understood multi-component transmembrane process^1,3^. At its core, simultaneous synthesis and secretion of individual cellulose chains is achieved by processive cellulose synthase (CesA) enzymes, which comprise a glycosyltransferase domain tightly associated with a channel-forming transmembrane domain^4^. CesAs utilise a UDP-glucose donor substrate to extend and then translocate the nascent polysaccharide’s cytosolic terminus, positioning it at the enzyme’s acceptor site before the onset of each catalytic cycle^5,6^.

Cellulose biosynthesis inhibitors (CBIs) are powerful tools in both research and agriculture^7,8^. For example, isoxaben (itself a commercially important herbicide, traditionally thought to directly target CesAs) and other CBIs are used extensively to study plant cell wall biosynthesis and cell wall integrity signalling. CBIs are often presumed to imitate the effects of CesA mutants and cell wall damage^8^. However, with no insights into any inhibition mechanism, and with recent reports questioning that isoxaben directly targets CesAs^9,10^, a molecular understanding of CBI activity is urgently required.

Thaxtomin A, a cyclic dipeptide, is the main phytotoxin of *Streptomyces* soil pathogens (such as *S. scabies*)^11,12^, which inflict major economic damage to world agriculture through various diseases of root and tuber crops^13–16^. The toxin exhibits an essential and highly unique 4-nitroindole pharmacophore, whose biosynthesis has been the subject of exceptional scientific interest^17–21^. However, while some data suggest thaxtomin A acts as a CBI^22–24^, its true molecular and pathological functions remain unknown. Indeed, its CBI effects have been proposed to be indirect^8^ and perhaps secondary to alternative activities, from immune system elicitation to mitochondrial toxicity^25–28^.

Here, we provide the first mechanistic description of a CBI from atomistic to organismal scales. Challenging long-existing dogma, *in vitro* assays demonstrate that most previously reported CBIs are poor inhibitors of purified CesAs, with the surprising exception of thaxtomin A. This compound targets CesAs and related cellulose synthase-like D (CslD) enzymes with unexpected directness, low-nanomolar affinity, and broad-spectrum activity. High-resolution cryo-EM structures of CesA bound to thaxtomin A show that the toxin occludes the enzyme’s cellulose secretion channel at its entrance’s highly conserved acceptor binding site, employing a self-locking mechanism and unique nitroaromatic electronic effects that target the enzyme’s catalytic centre. Based on these structures, we design partially resistant CesA mutants in an important step towards disease immunity. Finally, *in vivo* experiments in plants and algae establish the role of thaxtomin A in *Streptomyces* pathology and illuminate the osmotic and biochemical effects of a total arrest of cellulose biosynthesis, underscoring the fundamental role of cellulose in cell walls.

## Results

### Thaxtomin A specifically targets cellulose synthases in vitro

To our knowledge, no CBI has ever been tested in a chemically defined *in vitro* system. Thus, we attempted to detect inhibition of three previously characterised^4,29^ CesA isoforms purified in GDN detergent: CesA8 from hybrid aspen (*Ptt*CesA8; involved in secondary cell wall biosynthesis), and CesA1 and CesA3 from soybean (*Gm*CesA1 and *Gm*CesA3; involved in primary cell wall biosynthesis) (**Supplementary Fig. 1**). Of note, recombinantly expressed CesAs purify as monomeric and trimeric species, with the homotrimers likely representing physiologically relevant complexes. Because the *in vitro* catalytic activity of both species is comparable,^29^ both were used for activity measurements (see Methods).

Employing a radiolabelled substrate assay (in which CesA incorporates ^3^H-glucosyl moieties into ethanol-insoluble material), we designed an inhibitor screen using the following putative CBIs: isoxaben, thaxtomin A, dichlobenil, flupoxam, indaziflam, and acetobixan^8^, including 5 % DMSO to maintain solubility. At excess concentrations (0.5 mM), only isoxaben, thaxtomin A, and flupoxam showed apparent inhibition (**Fig. 1a**), with excess isoxaben also failing to inhibit *Gm*CesA1.

**Fig. 1.**
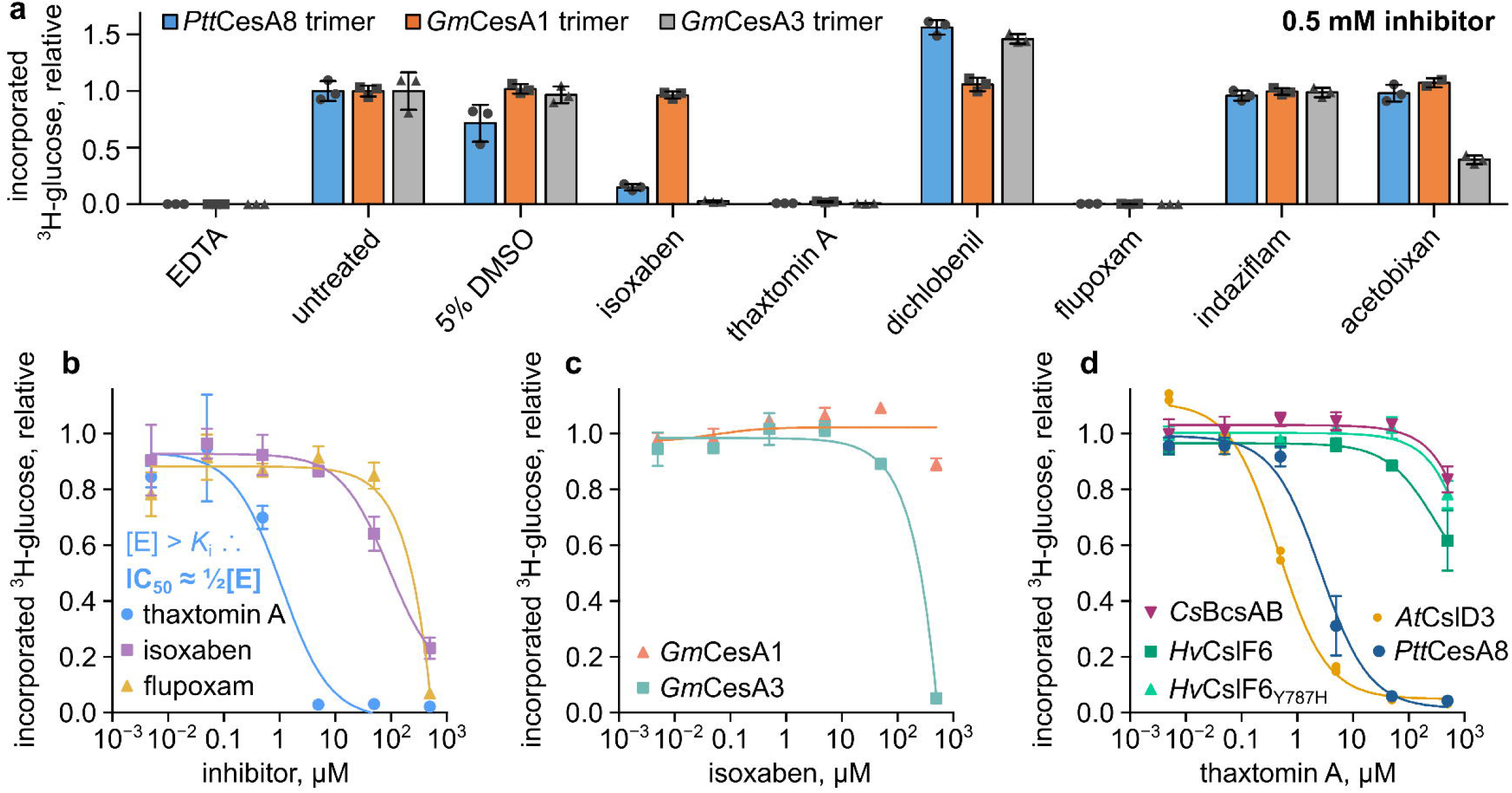
Out of various reported inhibitors, only thaxtomin A substantially inhibits cellulose synthase *in vitro*. Inhibitor screening by tritium incorporation assay. Purified cellulose synthase (CesA) isoforms and related enzymes were incubated overnight at 4 μM concentration with tritiated UDP-glucose in the presence of putative inhibitors (as well as the known inhibitor EDTA). Ethanol-insoluble products were quantified by scintillation counting. Means and SEM result from three technical replicates. **a** Initial screen at high inhibitor concentration (0.5 mM) in 5 % DMSO. Activities were normalized to the untreated control. **b** Inhibition of *Ptt*CesA8 by varying concentrations of isoxaben, flupoxam, and thaxtomin A in the presence of 5 % DMSO. Note that for thaxtomin A, the apparent IC_50_ is below enzyme concentration, thus IC_50_ > *K*_i_ (tight binding problem). **c** Inhibition (poor) of primary cell wall CesA isoforms by isoxaben. **d** Inhibition of *Ptt*CesA8, Arabidopsis CslD3, *Hv*CslF6, a β-1,4-glucan synthesizing *Hv*CslF6 mutant (Y787H; middle), and bacterial cellulose synthase (*Cs*BcsAB) by thaxtomin A (GDN-solubilized inhibitor). For **b** and **c**, average background activity seen in the EDTA control was subtracted from the other measurements, which are normalized to the uninhibited control. Low protein yields permitted only two technical replicates for *At*CslD3 (shown as individual data points).

However, when we tested these three putative inhibitors at lower and thus more physiological concentrations, these results were not reproduced. Indeed, despite their known *in vivo* toxicity at nanomolar concentrations^30,31^, neither isoxaben nor flupoxam showed any inhibitory effect at 5–50 μM concentration or lower (**Fig. 1b,c**). By contrast, thaxtomin A was able to efficiently titrate the activity of *Ptt*CesA8, with an apparent IC_50_ of ∼2 µM (**Fig. 1b**). However, with the concentration of CesA being 4 μM in these assays, the true binding constant is much lower than the IC_50_ due to the ‘tight binding problem’^32^, wherein the concentration of free inhibitor is substantially depleted by sequestration inside the enzyme. Thus, out of the six compounds, thaxtomin A was the only effective inhibitor of CesA.

Next, to eliminate potential negative effects of DMSO, we solubilised thaxtomin A in GDN detergent (see Methods). Subsequently, we tested this material on other CesA-related enzymes purified in GDN, namely Arabidopsis β-1,4-glucan synthase CslD3 (*At*CslD3) and barley mixed-linkage glucan (MLG) synthase CslF6 (*Hv*CslF6) (**Fig. 1d**). CslDs are thought to constitute non-canonical cellulose synthases, while *Hv*CslF6 produces a glucan with both β-1,4 and β-1,3 linkages^33^. Interestingly, *At*CslD3 was also inhibited by thaxtomin A, whereas the closely related *Hv*CslF6 showed substantial resistance. However, this was not due to their differing polysaccharide products, as the previously characterised *Hv*CslF6[Y787H] mutant, which produces mainly β-1,4-glucan, also exhibited similar resistance. Finally, the purified bacterial BcsAB cellulose synthase complex from *Cereibacter* (*Rhodobacter*) *sphaeroides* (**Supplementary Fig. 1**) also exhibited resistance to thaxtomin A inhibition, consistent with the evolutionary divergence of plant and bacterial CesAs.

### Thaxtomins block CesA’s channel entrance

To confirm that thaxtomin A binds directly to CesA, we employed nano differential scanning fluorimetry (nanoDSF), which detects changes to the protein’s thermal unfolding profile due to ligand binding. Indeed, indicative of tight binding, stoichiometric amounts of thaxtomin A significantly stabilised *Ptt*CesA8, raising the temperature of inflection (*T*_i_; related to melting point, *T*_m_) from 60 °C to 72 °C (**Extended Data Fig. 1**).

To resolve how thaxtomin A inhibits *Ptt*CesA8, we employed single-particle cryo-EM. However, standard cryo-EM approaches have so far only achieved CesA maps of limited resolution (generally > 3.0 Å), with weaker and discontinuous density in the transmembrane (TM) regions^4,29,34^, which can obscure molecular details such as small-molecule ligands. To resolve this, we developed an advanced data processing pipeline (**Supplementary Fig. 2**) to obtain high-resolution structural information for CesA. Most critically, we found that a symmetry expansion approach was necessary to overcome the symmetry-breaking flexibility of the CesA trimers. This permitted improved alignment and thus overall map quality, especially in the transmembrane region (**Supplementary Fig. 3**). Our combined strategy ultimately achieved an estimated resolution of 2.2 Å. While being consistent overall with the previously generated model,^4^ our new map reveals important new molecular insights, including water coordination at the active site and rotamer-level details of the TM region.

Most importantly, close inspection of the map (**Fig. 2a**) revealed the presence of a strong non-proteinaceous density at the acceptor site of CesA’s TM channel. This density was present in all three protomers and perfectly accommodated a thaxtomin A molecule (**Fig. 2b,c, Supplementary Fig. 3**). Strikingly, thaxtomin A’s unique nitroindole group stacks directly with Trp718 of the characteristic QxxRW motif (**Fig. 2d,e**), which sits at the cytosolic entrance to CesA’s cellulose secretion channel and normally positions the acceptor glucosyl moiety of the nascent cellulose chain. With interactions between tryptophan and nitroaromatic groups already known to be exceptionally strong^35^, this contact appears to be a key function of the nitroindole group.

**Fig. 2.**
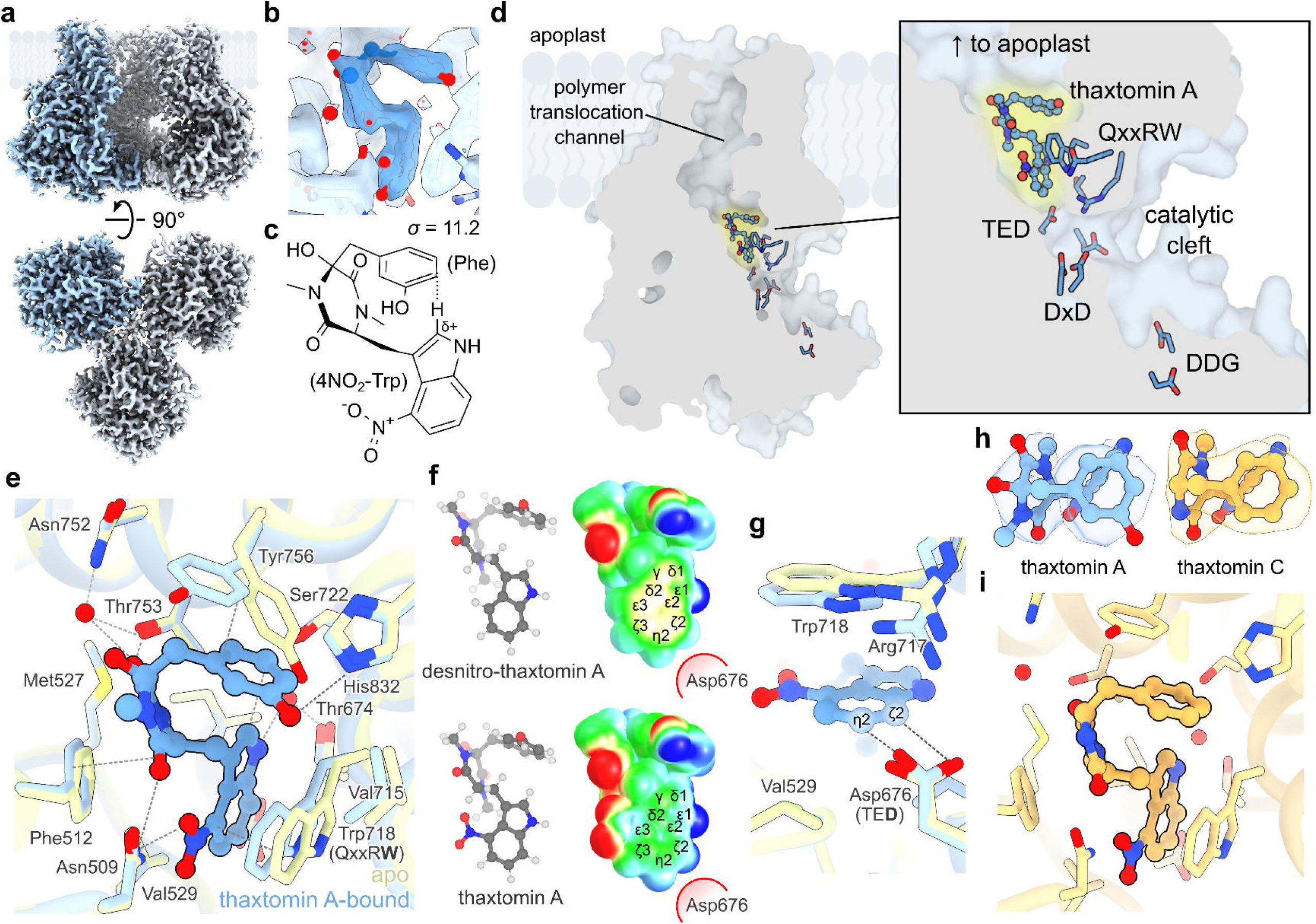
Thaxtomin utilizes its unusual chemistry to bind to the transmembrane channel entrance. **a** Overall cryo-EM density map for *Ptt*CesA8 bound to thaxtomin A (composite map of symmetry-expanded monomers). **b** Local cryo-EM density for the thaxtomin A ligand. **c** Structural formula for thaxtomin A, showing folded conformation. **d** Location of thaxtomin A binding site in the transmembrane channel of *Ptt*CesA8 (single protomer shown). Important sequence motifs (DDG, DxD, TED, and QxxRW) are labelled. **e** Thaxtomin A binding site in detail. Pale blue: thaxtomin A-bound *Ptt*CesA8. Lime-green: apo *Ptt*CesA8. **f** Electrostatic potential maps for thaxtomin A with and without its nitro group. Blue = positive (+30 kcal/mol), red = negative (−30 kcal/mol). Note polarization of C^ζ2^ and C^η2^. **g** Thaxtomin A binding site shown from the perspective of the catalytic base (Asp676). **h** Local cryo-EM density (σ = 11.2) for thaxtomin A vs thaxtomin C, showing differences in density. **i** Thaxtomin C binding site. Waters shown as red spheres.

Two other important interactions with aromatic side chains are notable: firstly, amide–π stacking between the central diketopiperazine ring of thaxtomin A and Phe512, and secondly, π-stacking between thaxtomin A’s phenol group and Tyr756. The latter results from the reorientation of Tyr756’s side chain towards the apoplast, partially occluding the channel. Diffusion of thaxtomin A towards the extracellular side is therefore hindered, ‘locking’ itself in place.

In addition to stacking with Trp718, the unusual nitroindole group directly contacts Asp676, CesA’s base catalyst and the final residue of the TED motif. The strongly electron-withdrawing nitro group imparts a partial positive polarisation upon the C^ζ2^ and C^η2^ carbons on the opposite side of the nitroindole ring system (**Fig. 2f**), and intriguingly, these carbon atoms directly abut Asp676’s anionic carboxylate group (**Fig. 2g**). In a cellulose-bound state, this residue forms a strong H-bond with the acceptor glucose’s C4 hydroxyl group in order to deprotonate it during catalysis^5,36^. Thus, we probed whether thaxtomin A’s nitroindole group mimics the acceptor by forming unusual C–H···O interactions with the catalytic base. Indeed, modelled as 4-nitroindole and acetate, quantum chemistry calculations estimate an interaction energy of 5.17 kcal/mol for a local dielectric constant (ε) of 4 (protein interior) or 2.11 kcal/mol for ε = 78 (i.e. water), which is comparable to a typical H-bond interaction, as calculated for a dimer of formamide (4.90 or 3.79 kcal/mol, respectively). Thus, *Streptomyces* pathogens’ bespoke nitroaromatic chemistry serves to form unique and unprecedented biochemical interactions with CesA’s catalytic base and the surrounding acceptor site.

In addition, thaxtomin A binding is supported by several hydrogen bonds (H-bonds) with *Ptt*CesA8, including between Asn509 and the 4-nitrotryptophan ‘backbone’ carbonyl (and/or the nitro group) and Thr753 and the phenylalanine backbone carbonyl (**Fig. 2e**). Our structure also indicates an H-bond between thaxtomin A’s phenol hydroxyl and His832, though the converse phenol orientation permitting H-bonding with Ser722 is not fully ruled out. Interestingly, two well-ordered water molecules also appear to participate in thaxtomin A binding: the first coordinated by Thr674 and Ser722 and the second by Asn752 and Thr753 (**Fig. 2e, Supplementary Fig. 4**). Each forms a canonical H-bond with the nitroindole amine and the phenylalanine backbone carbonyl, respectively.

Thaxtomin C, the main toxin of *S. ipomoeae* (responsible for sweet potato soil rot)^37^, is a congener of thaxtomin A that lacks the latter’s phenylalanine *N*-methyl group and hydroxyl decorations (**Fig. 2c,h**). To test whether thaxtomin C also binds to CesA, we resolved its structure bound to *Ptt*CesA8, ultimately producing a 2.5 Å-resolution map (**Supplementary Fig. 5**). Indeed, thaxtomin C’s distinct density (**Fig. 2h, Supplementary Fig. 6**) adopts the same binding pose and interactions as thaxtomin A, except that, importantly, it lacks H-bonding with His832 (**Fig. 2i**). Interestingly, the S-shaped conformation of both thaxtomins, which is likely stabilised by intramolecular C–H···π stacking (**Fig. 2c**), mirrors a previous X-ray crystal structure of the thaxtomin A compound itself^38^, as well as resembling the conformations of thaxtomins B and D during biosynthetic hydroxylation^39^. Thus, thaxtomins may exhibit preferred conformations or perhaps even stable secondary structures, reinforcing their shape complementarity to the CesA binding site.

### High-affinity non-competitive inhibition

To obtain kinetic parameters for CesA’s inhibition, we overcame the tight binding problem through the use of the UDP-Glo method^40^, which converts free UDP reaction byproducts into luminescence, and whose high sensitivity could detect activity from as little as 1 nM *Ptt*CesA8. Collecting Michaelis-Menten data under different inhibitor concentrations, we observed that *V*_max_ was substantially reduced by the presence of inhibitor, whereas the *K*_M_ for UDP-glucose was only mildly affected (**Extended Data Fig. 2**). Though these characteristics are typical for tight-binding inhibitors^32^, they are consistent with a non-competitive or quasi-non-competitive inhibition mechanism with respect to UDP-glucose.

Subsequent experiments to determine thaxtomin A’s apparent inhibition constant (*K*_i_*) using the tight inhibition equation of Copeland^32^ (see Methods) produced a highly reproducible and remarkably low estimated *K*_i_* of 2.1 nM (**Fig. 3a**). Note that, having ruled out an ‘uncompetitive’-type inhibition mechanism in which the inhibitor binds to the enzyme–substrate complex, *K*_i_* forms the upper bound for true *K*_i_. By contrast, thaxtomin C’s *K*_i_* was substantially higher, at 49 nM (**Fig. 3b**), likely due to the lack of the H-bond with His832 (**Fig. 2i**). Furthermore, as observed in the tritiated glucose incorporation assay (**Fig. 1d**), *Hv*CslF6 was substantially resistant to inhibition by thaxtomin A (*K*_i_* = 14 μM; **Fig. 3b**).

**Fig. 3.**
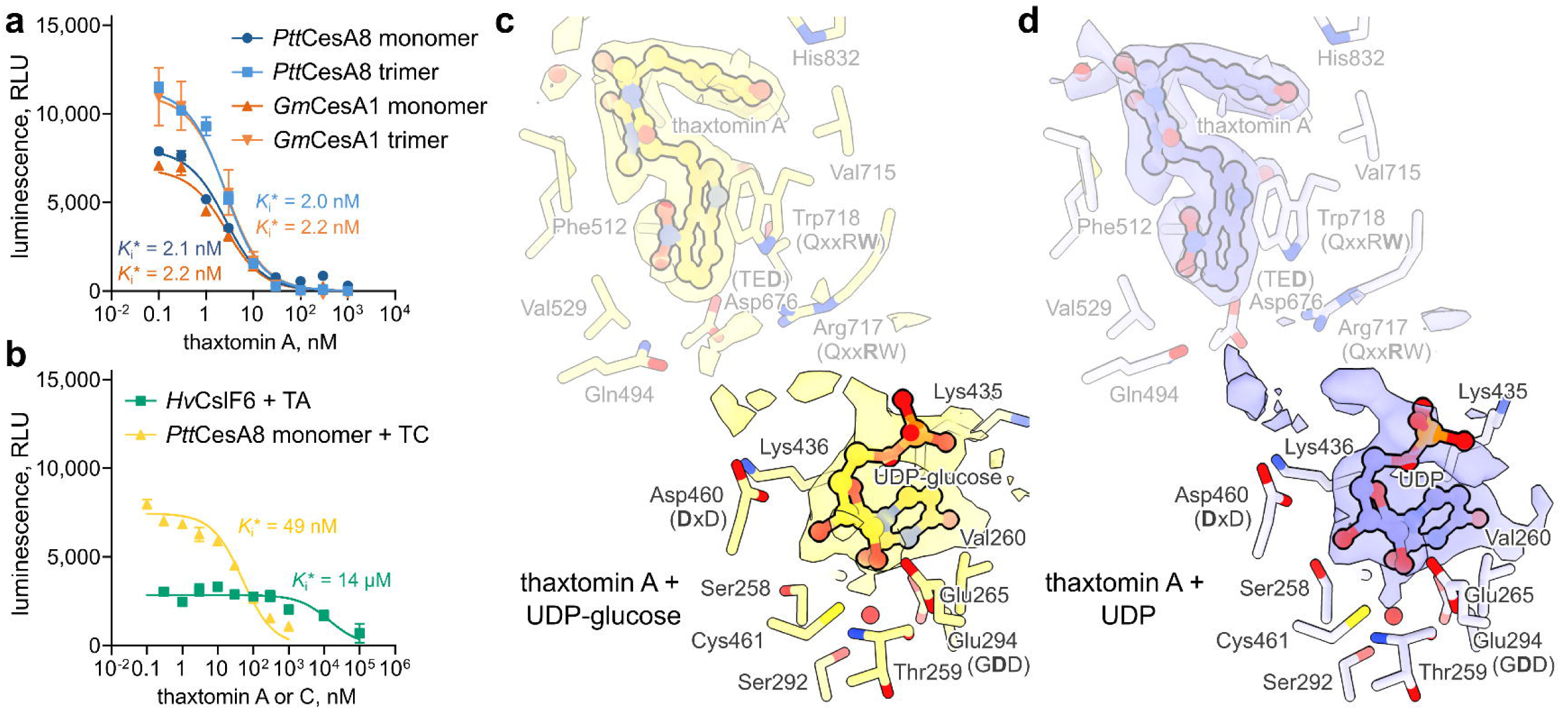
Thaxtomin A binds tightly to cellulose synthase with low-nanomolar affinity and hinders substrate insertion. **a** Inhibition of *Ptt*CesA8 activity as observed by the UDP-Glo assay. Enzyme (1 nM) was pre-incubated with 5 % DMSO ± thaxtomin A for 2 h before incubation with UDP-glucose substrate for 1 h at 30 °C. Free UDP is converted into luminescence. *K*_i_*: apparent inhibition constant (*K*_i_* ≥ *K*_i_). **b** Inhibition of *Ptt*CesA8 by thaxtomin C (TC) and *Hv*CslF6 by thaxtomin A (TA). For **a** and **b**, curves are fitted with the modified Morrison equation, and each mean and standard error results from six replicates, comprising two full biological replicates (separate protein purification and assay) each with three technical replicates. **c, d** Cryo-EM structures of *Ptt*CesA8 incubated with thaxtomin A, MgCl_2_, and either UDP-Glc (c) or UDP (d). The catalytic pocket and transmembrane channel are shown in surface representation, while experimental cryo-EM density for thaxtomin A and UDP(-Glc) is shown at a low contour level (σ = 4.0). Due to ambiguous densities, neither glucosyl nor β-phosphate residues were modeled. Waters shown as red spheres.

To confirm the non-competitive inhibition mechanism, we solved further high-resolution structures of *Ptt*CesA8 in complex with thaxtomin A, Mg^2+^, and UDP or UDP-glucose (**Supplementary Figs. 7 and 8**). In both structures, densities corresponding to the uracil, ribose, and α-phosphate moieties of UDP/UDP-glucose were clearly resolved (**Fig. 3c,d, Supplementary Fig. 9**), demonstrating the ability of donor substrate to bind in the presence of excess thaxtomin A. Further, our high-resolution map highlights the role of a previously unknown water molecule, coordinated by Ser292 and the backbone of Thr259, in binding the substrate’s uracil moiety (**Fig. 3c,d**). However, at least in contrast to previous moderate-resolution structures of *Ptt*CesA8 in complex with UDP-glucose^6^, the substrate’s glucosyl and β-phosphate moieties were not well resolved, suggesting that these portions were disordered, despite the absence of any obvious steric clash (**Supplementary Fig. 10**). Combined, these data show that thaxtomin A blocks the transfer of glucosyl moieties from UDP to the enzyme’s acceptor site.

### Design of resistance mutants

Thaxtomin-resistant CesA mutants would be a powerful tool in developing disease resistance as well as engineering selective herbicides. However, the strict residue conservation of the thaxtomin binding site among CesA isoforms (**Supplementary Fig. 11**), which presumably arises from its role in coordinating the glucosyl acceptor during catalysis, likely prevents the natural development of toxin resistance. Indeed, even *At*CslD3 exhibits only relatively minor differences at this site, namely the substitution of Thr753 and Ile677 in *Ptt*CesA8 for valines in *At*CslD3 (**Fig. 4a**), explaining why thaxtomin A inhibits both enzymes. On the other hand, *Hv*CslF6, which is resistant to thaxtomin A, exhibits two other differences in the binding site: the substitution of His832 for tyrosine (Tyr787 in *Hv*CslF6) and Thr753 for isoleucine (Ile708)^41^. In both cases, an H-bond partner residue is replaced by a bulky side chain that would sterically hinder thaxtomin A binding. Finally, in resistant *Cs*BcsAB, Pro757 of *Ptt*CesA8 is replaced with tryptophan (Trp417 in *Cs*BcsA), which occludes the phenol-binding subsite (**Fig. 4b**), consistent with the lack of sensitivity to thaxtomin A.

**Fig. 4.**
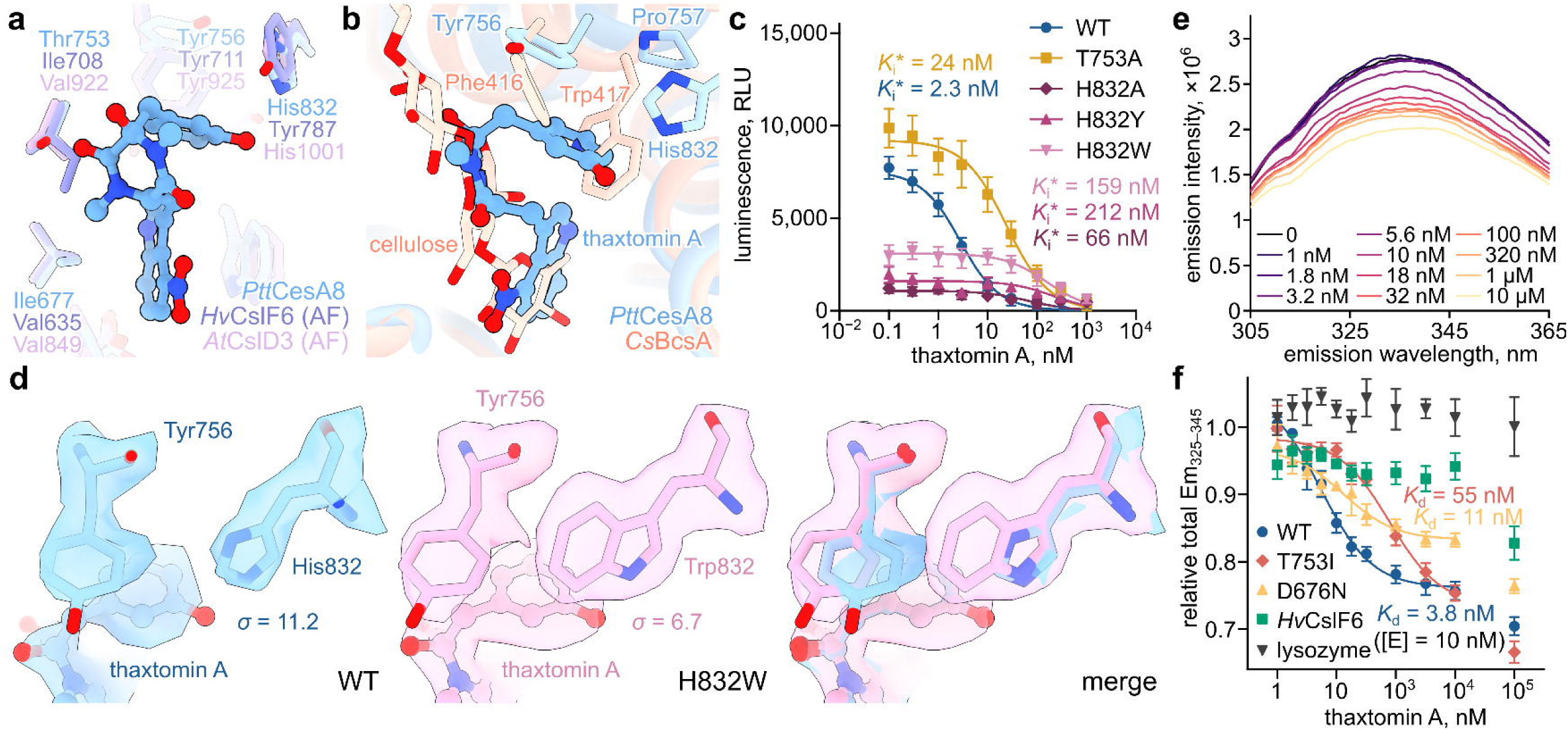
Point mutations to the thaxtomin binding site introduce partial resistance. **a,b** *Ptt*CesA8 thaxtomin A binding site aligned with AlphaFold predictions for *Hv*CslF6 and *At*CslD3 (a) or the experimental structure of *Cs*BcsA (b, PDB: 4P02), highlighting differing residues. **c** Activities and inhibition of the various mutants as measured by UDP-Glo. Means and SEMs result from two full biological replicates, each with three technical replicates. **d** Experimental cryo-EM structure of the *Ptt*CesA8[H832W] mutant bound to thaxtomin A and aligned with wild type, showing rearrangement of Tyr756 required to bind the inhibitor. **e** Tryptophan fluorescence quenching of *Ptt*CesA8 by thaxtomin A in the presence of 5 % DMSO. Enzyme and inhibitor were pre-incubated for 2 h at room temperature. Excitation wavelength: 283 nm. **f** Tryptophan fluorescence quenching of various enzymes and mutants by thaxtomin A (three independent technical replicates).

This considered, we attempted to design thaxtomin resistance mutations without perturbing CesA’s catalytic activity, ultimately purifying the following mutants: T753A, T753I, H832A, H832W, and the previously reported H832Y mutant^41^ (P757W and P757I mutants were attempted but failed to purify) (**Supplementary Fig. 12a**). Additionally, we purified a D676N catalytic base mutant to confirm the theoretical interaction between the nitroindole ring and the catalytic base (**Fig. 2g**).

We then used the UDP-Glo assay to monitor the inhibition of all but two of the CesA mutants (D676N and T753I), whose activities were below the detection threshold; **Supplementary Fig. 12b**). In all cases, thaxtomin A’s *K*_i_* was increased relative to wild-type enzyme, with H832Y exhibiting the largest increase to 212 nM (∼100-fold), albeit at the expense of overall catalytic activity **Fig. 4c**). The T753A mutant, however, exhibited a tenfold increase in *K*_i_* from 2.3 nM to 24 nM without any apparent decrease in activity.

Though the H832W mutant’s *K*_i_* was increased to 159 nM, we had anticipated this mutant’s resistance to be stronger, having predicted a steric clash between the introduced tryptophan and Tyr756’s reoriented position. A high-resolution structure of this mutant bound to thaxtomin A revealed that, despite inhibitor binding, the introduced Trp forces Tyr756 to adopt an unfavourable conformation with less efficient stacking with thaxtomin A’s phenol ring (**Fig. 4d, Supplementary Fig. 13**). Combined with the fact that Trp cannot act as an H-bond acceptor, these factors help to explain this mutant’s partial resistance.

Finally, making use of 4-nitroindole’s powerful fluorescence-quenching properties^42,43^, we established a Trp fluorescence quenching assay to determine the binding affinities of the two least-active mutants (T753I and D676N) for thaxtomin A (requiring 10 nM enzyme for adequate signal). Indeed, binding of thaxtomin A to *Ptt*CesA8 produced an appreciable drop in total Trp fluorescence (**Fig. 4e,f**). Because higher concentrations appeared to produce non-specific effects, we fitted the Copeland equation to data in the 0.001–1 µM range, producing an estimated binding constant of 3.8 nM for the WT enzyme. By contrast, the binding constants for the D676N and T753I were raised approximately three- (11 nM) and tenfold (55 nM), respectively, verifying the importance of nitroindole electrostatic interactions and suggesting directions for further mutagenesis.

### Thaxtomins target expanding cells in vivo

While thaxtomin A’s ability to inhibit CesA is consistent with its previously observed effects on seedling morphology, cell wall composition, and CesA dynamics^22–24^, a significant body of literature suggests that the toxin could achieve cell death through other pathways. Furthermore, the surprising ineffectiveness of canonical CBIs to inhibit CesA *in vitro* challenges important previous interpretations based on their presumed activities. Therefore, having identified thaxtomin A as the so far only confirmed *bona fide* CBI, we sought to better characterise its effects *in vivo* to assess the true consequences of a total arrest of cellulose biosynthesis.

Firstly, we attempted to identify secondary targets of thaxtomin A by comparing its effects on cellulosic and non-cellulosic land plant-adjacent algal species. Accordingly, we studied toxicity against four streptophyte algae: *Closterium peracerosum-strigosum-littorale* complex, with a highly cellulose-rich wall;^44,45^ *Mesostigma viride*, which appears to lack cellulose entirely^46,47^; and *Klebsormidium nitens* and *Chlorokybus riethii*, which exhibit cellulose as a minor component^48–50^.

Notably, and in contrast to isoxaben, thaxtomin A exhibited acute toxicity towards *Closterium p.s.l.* (**Fig. 5a**). Lower doses (300 nM) induced cell wall bulging at the isthmus (the site of wall expansion) whereas higher concentrations disintegrated cells (**Fig. 5b**). By contrast, its effects on the other three species were substantially milder or undetectable (by quantifying chlorophyll fluorescence or absorbance), with a minor effect on *M. viride* growth perhaps attributable to the yellow-coloured compound’s attenuation of photosynthetic frequencies (**Fig. 5a,c–e**). Overall, sensitivity to thaxtomin A appeared to correlate strongly with the importance of cellulose to the cell wall integrity of each organism.

**Fig. 5.**
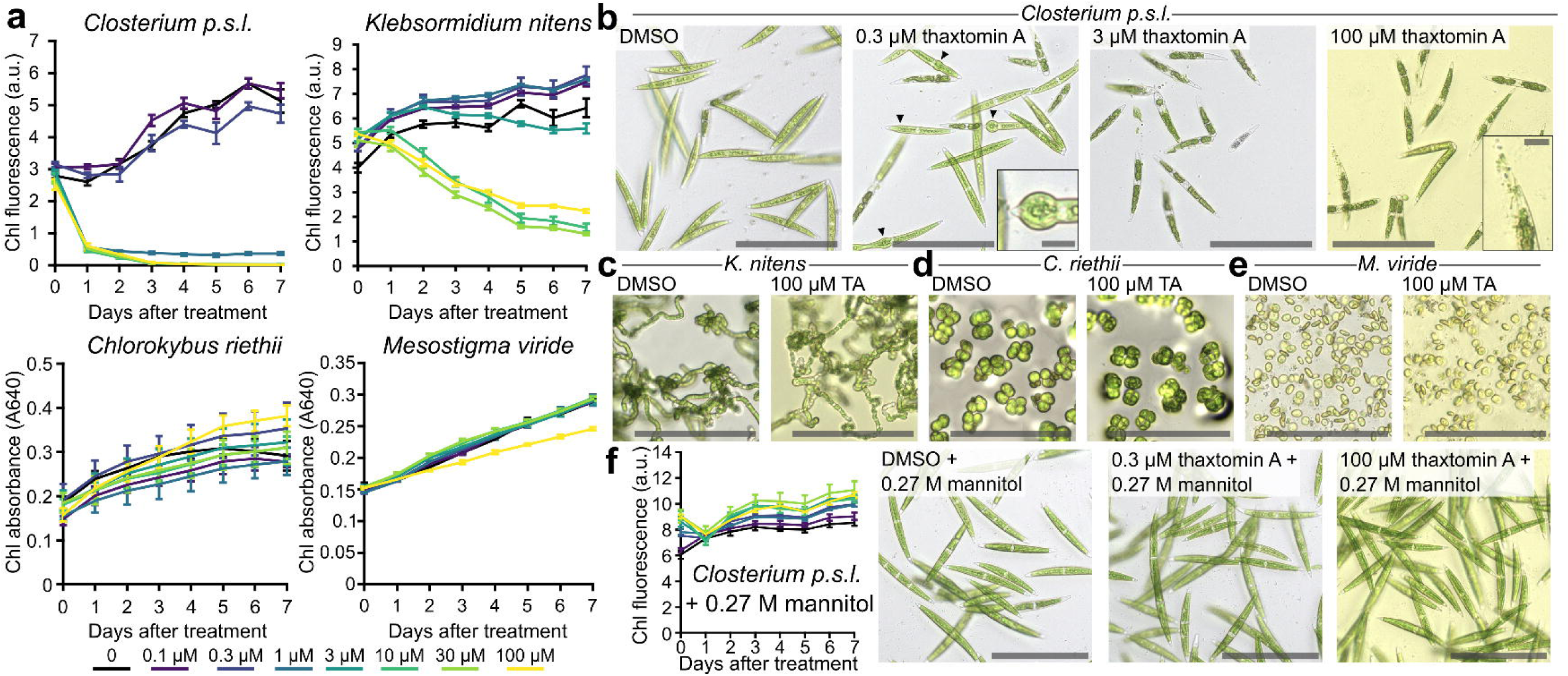
Thaxtomin A induces osmotic lysis of cellulose-dependent streptophyte algae. Streptophyte algae *Closterium peracerosum-strigosum-littorale* (high cellulose content), *Klebsormidium nitens* (low cellulose content), *Chlorokybus riethii* (very low cellulose content), and *Mesostigma viride* (no cellulose) were grown in the presence of 0.5 % DMSO and 0–100 μM thaxtomin A. **a** Growth curves measured using chlorophyll fluorescence (*Closterium p.s.l.* and *K. nitens*) or absorbance (*C. riethii* and *M. viride*). Error bars: standard error of the mean (SEM) of three technical replicates. **b** *Closterium p.s.l.* cells after three days in the presence of 0–100 μM thaxtomin A. Cell wall bulges are labelled with arrowheads. Scale bars: 150 μm (inset: 15 μm). **c, d, e** Morphology of *K. nitens*, *C. riethii*, and *M. viride*, respectively, after seven days in the presence of 0 or 100 μM thaxtomin A. Scale bars: 150 μm. **f** Growth of *Closterium p.s.l.* treated with thaxtomin A in the presence of 0.27 M mannitol. Cells were imaged after three days.

Remarkably, we were able to fully ameliorate these toxic effects by osmotic protection with 0.27 M mannitol^51^. (**Fig. 5f**). This suggests that thaxtomin-induced cell death is primarily due to osmotic lysis resulting from cellulose depletion and a consequent loss of cell wall integrity. These results not only demonstrate thaxtomin A’s supremely broad specificity (extending to algal CesAs) but also validate the fundamental cell wall turgor hypothesis that cellulose microfibrils constrain osmotic forces in plants.

To investigate thaxtomin A’s effects on vascular plants, we employed the model plant *Nicotiana benthamiana* (closely related to important *Streptomyces* hosts such as potato). Though previous studies have reported that thaxtomin A can induce a hypersensitive immune response, Ca^2+^ signalling, and the secretion of cellotriose, our own leaf inoculation assays, Arabidopsis apoaequorin Ca^2+^ reporter experiments, and carbohydrate electrophoresis analyses of *N. benthamiana* seedling exudates do not support these ideas (see Methods and **Supplementary Figs. 14 – 16**).

Furthermore, almost all previous studies on the effects of thaxtomins have examined seedlings grown in the constant presence of toxin^22–24^, whose altered morphology could result from developmental effects originating shortly after germination. To better mimic *in vivo* conditions during root infection by *Streptomyces* spp., we permitted *N. benthamiana* seedlings to develop normally for nine days before transplanting to medium with DMSO, 10 μM thaxtomin A, or 10 μM isoxaben. Indeed, we noticed several striking differences in root tip morphology under brightfield and confocal microscopy (the latter employing calcofluor white staining of epidermal cell walls). Compared with DMSO, thaxtomin A treatment arrested cell elongation, induced cell bulging, and evoked a dark-brown discolouration (**Fig. 6a,b, Supplementary Video 1**), with similar results observed in five-day-old rice seedlings (**Supplementary Fig. 17**). Furthermore, contrasting previous studies using constant toxin exposure, these effects were tightly localised to the root elongation zone where cells are actively expanding and synthesising new cell wall.

**Fig. 6.**
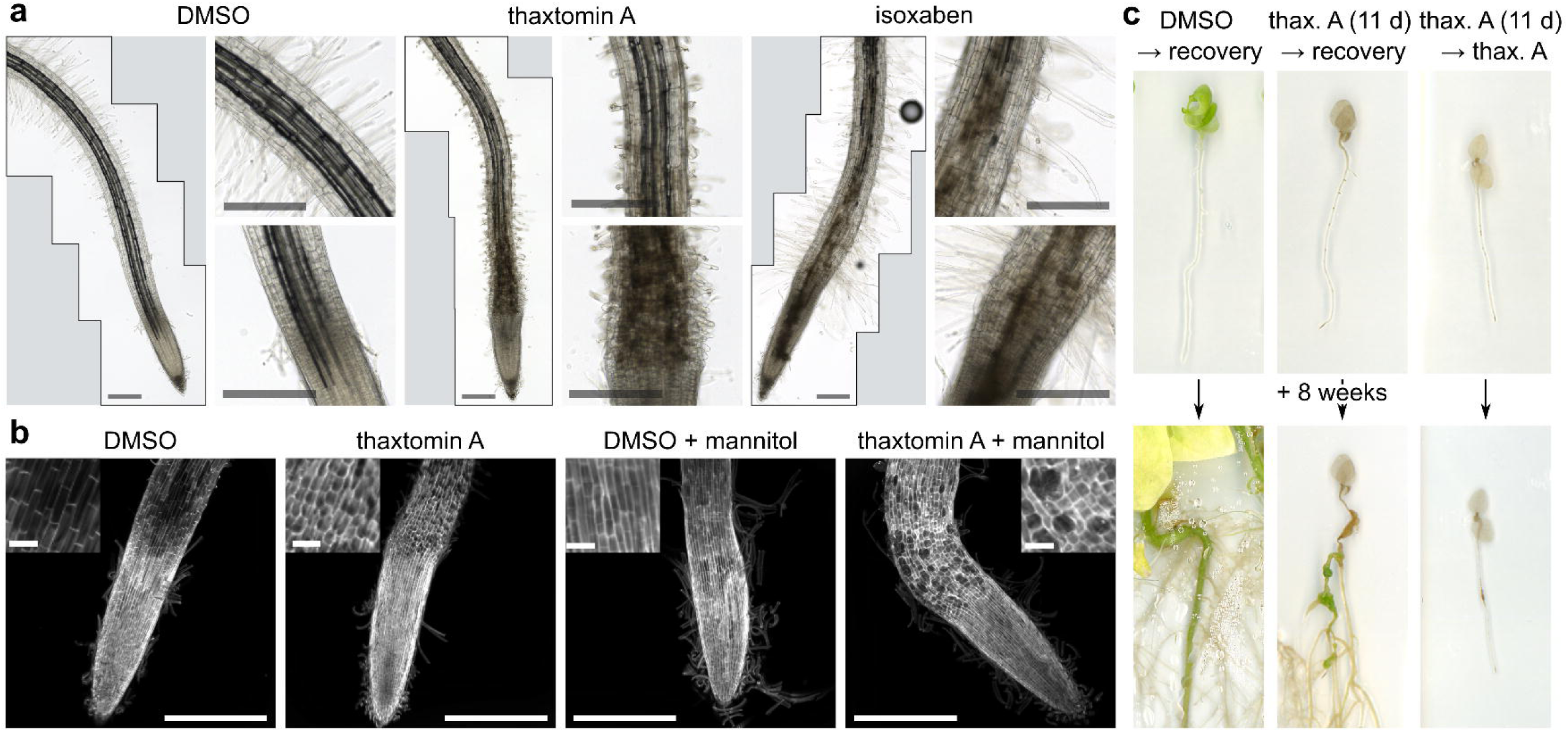
Thaxtomin A specifically disrupts root elongation zone morphology in an osmotic stress-dependent manner. *Nicotiana benthamiana* seedlings were grown for nine days under normal conditions before treatment with 10 μM thaxtomin A or isoxaben. **a** Short-term (24 h) effects on root hairs and elongation zone (representative of ten biological replicates). Scale bars: 300 μm. **b** Short term effects of thaxtomin A and/or 0.3 M mannitol treatment on representative root tips, as visualised by calcofluor white staining and confocal microscopy. Scale bars: 500 μm (inset: 50 μm). **c** Recovery from long-term thaxtomin A treatment. Nine-day-old *N. benthamiana* seedlings were treated with DMSO/thaxtomin A for 11 days before transplanting to solid medium lacking (or retaining) thaxtomin A and observed for eight weeks (representative plants from **Supplementary Fig. 18**).

Moreover, while isoxaben mirrored some of these effects (albeit in a milder and less specifically localised manner), its effects on root hairs contrasted those of thaxtomin A (**Fig. 6a**). Similar to previous results in Arabidopsis, isoxaben had very little effect aside from an occasional ‘ballooned’ appearance. Conversely, thaxtomin A treatment severely stunted the growth of new root hairs. This is consistent with our discovery that thaxtomin A inhibits CslDs, which are thought to be responsible for synthesising cellulose in root hair cell walls.^52^

Unexpectedly however, and in contrast to its effects on algae, increasing external osmolarity with 0.3 M mannitol exacerbated the effects of thaxtomin A, resulting in severe swelling (**Fig. 6b, Extended Data Fig. 3**). While the molecular basis for this effect remains to be determined, we speculate that stress-induced signalling, cell wall changes, and water influx from neighbouring cells may be primarily responsible.

Finally, we found that young seedlings have an astonishing ability to recover after 11 days treatment with thaxtomin A. Indeed, within two weeks of transplantation to recovery medium, formerly thaxtomin-treated seedlings established new lateral roots, eventually developing a substantial root system as well as unusual photosynthetic root galls (the aerial tissues having been lost presumably due to water stress; **Fig. 6c, Supplementary Fig. 18**). This demonstrates not only the limits of thaxtomin A’s cellular toxicity but shows that unexpanding plant cells exhibit stable viability in the absence of cellulose biosynthesis.

## Discussion

Here, we have identified thaxtomin A as the first confirmed *bona fide* CBI, demonstrating its unique directness, avid affinity, and breadth of targets. This necessitates a paradigm shift in the use and interpretation of CBIs in research, indicating that thaxtomin A, not isoxaben or dichlobenil, should be considered the canonical CBI. Nevertheless, we cannot rule out the possibility that the latter compounds achieve their *in vivo* effects by perturbing higher-order CesA complexes^53,54^, which could potentially relate to the isoform-specific effect we observed for excess isoxaben. However, known differences between isoxaben and thaxtomin A’s biological effects (such as in ectopic lignification and their relationship to the THESEUS1 receptor^24,55^) warrant a re-evaluation of existing results and experimental practices.

Through the process of bacterial evolution, thaxtomins have been precisely tailored to CesA’s acceptor site, succeeding in what manmade herbicides have apparently only aspired to. Their basic functionality is provided by the unique chemistry of the nitroindole group and the shape of their cyclic dipeptide scaffold, further honed by additional methyl and hydroxyl decorations. Thus, these molecules provide an elegant masterclass in targeting the semi-hydrophobic TM channel of CesA and related enzymes. Interestingly, two important components of the nitroindole binding site (the QxxRW motif and catalytic x(E/D)D motif) are exceptionally well conserved in related synthases of polysaccharides such as chitin, hyaluronan, and bacterial cellulose^56,57^, presenting this pharmacophore as a new and promising seed for the generation of novel herbicides, antibiotics, antifungal agents, and anticancer drugs.

Our results shed insight into how *Streptomyces* pathogens have subverted their originally symbiotic niche into a parasitic one, evolving from a saprotrophic diet of discarded cellulosic material to attack their erstwhile partners^13,58^. Thaxtomin A apparently allows these pathogens to weaken the cell walls of expanding host cells for hyphal invasion, with toxic osmotic and immune effects as secondary consequences. Furthermore, the high sequence conservation of the thaxtomin binding site ensures broad specificity and suppresses natural resistance. Nevertheless, we show that resistance can in principle be designed without deleterious side-effects, raising hope for genetically engineered immunity against these economically important plant diseases.

Finally, we provide some of the first mechanistically qualified evidence for the fundamental functions of cellulose in plant biology. Thus, thaxtomin A presents itself as an indispensable tool for arresting, modulating, and controlling total cellulose biosynthesis in plants.

## Methods

### Plant growth

Except where stated below, algae and seedlings were grown under long-day conditions (16 h photoperiod) under 10 μmol m^−2^ s^−1^ (algae) or 80 μmol m^−2^ s^−1^ (plants) photosynthetic photon flux at 23 °C in a Percival Scientific AL36L4C8 growth chamber equipped with SciBrite LED lights.

*N. benthamiana* seeds were a gift of Prof. Michael Timko, University of Virginia. Rice (*Oryza sativa* ssp. *japonica* ‘Kitaake’) seeds were a gift of Prof. Nam-Chon Paek, Seoul National University. For sterile plant cell culture, *N. benthamiana* seeds were surface-sterilised with 70 % ethanol for 20 s followed by 50 % household bleach (3 % hypochlorite final concentration) for 7 min, with shaking. After five washes with sterile water, seeds were sustained on half-strength Murashige and Skoog (½ MS) medium: 0.22 % Murashige and Skoog basal salts (#M5524, Sigma-Aldrich, Burlington, MA, USA), 1 % sucrose, 0.1 % MES-KOH, pH 5.7, and, for solid medium, 0.8 % plant tissue culture agar. Liquid *N. benthamiana* cultures were grown at 21 °C with shaking at 100 rpm under constant illumination from a T5HO 6500K fluorescent lamp (Philips) providing 80 μmol m^−2^ s^−1^ photosynthetic photon flux.

For leaf toxicity assays, *N. benthamiana* seeds were sown on germination mix (Pro-Mix FPX). Two weeks later, seedlings were transplanted to Lambert LM-111 all-purpose mix in ∼500-cm^3^ pots in the University of Virginia Department of Biology greenhouse under long-day conditions with 300–400 μmol m^−2^ s^−1^ illumination, 26–29 °C day temperature, 24–27 °C night temperature, and 30–36 % humidity. Plants were fertilised by constant liquid feed with 15-5-15 fertiliser at 150 ppm nitrogen.

For Ca^+2^ response assays, Arabidopsis seeds were surfacelzlsterilised with 2% Triton/70% ethanol and subsequently stratified for 3 days in the dark at 4°C. Seedlings were then grown in 96lzlwell plates (one seedling per well) under a 14lzlh photoperiod in liquid ½ MS supplemented with 1% sucrose at 19–22=°C.

*C. peracerosum-strigosum-littorale* complex NIES-67, *K. nitens* NIES-2285, *C. riethii* NIES-160, and *M. viride* NIES-296 were provided by the National Institute of Environmental Studies, Japan, through the MEXT National BioResource Project. Algae were grown in C medium^59^ and subcultured every 2–3 weeks. For *C. riethii*, colonies were maintained on C medium supplemented with 1.5 % plant tissue culture agar, whereas *C. riethii* thaxtomin toxicity assays were conducted in C medium supplemented with 0.4 % Phytagel gellan gum and 3 mM CaCl_2_.

### Algal toxicity assays

Algae were grown over the course of a week in the presence of 0.5 % DMSO and 0–100 μM thaxtomin A (SML1456; Sigma-Aldrich). Liquid cultures in mid/late log phase were diluted two- to three-fold in fresh medium and split into 24 individual cultures (three per concentration of inhibitor). For *Closterium p.s.l.* and *K. nitens*, 300-μL cultures (at 3×10^4^ cells mL^−1^ for *Closterium p.s.l.*) were grown in a SensoPlate glass-bottomed 96-well plate (Greiner Bio-One, Kremsmünster, Austria), with cells growing across the bottom of each well. To measure chlorophyll fluorescence, the bottom of the plate was scanned daily using an EVOS M7000 imaging system (ThermoFisher Scientific, Waltham, MA, USA) at 4X magnification using the Cy5 filter cube (Ex: 635 nm, Em: 692 nm). Additional brightfield images were acquired at 20X magnification. Total fluorescence signal for each well image was then obtained using ImageJ.

This approach was not possible for *M. viride* due to swimming behavior. Instead, 200-μL cultures at 2×10^6^ cells mL^−1^ were grown in a clear-bottomed, clear-walled 96-well plate and chlorophyll absorbance at 640 nm was measured daily using a SpectraMax i3x plate reader. Additional brightfield images were acquired for cultures grown in parallel in glass-bottomed plates. Similarly, a specialised approach was required for *C. riethii* as, for unknown reasons, we were unable to propagate this species in liquid medium. Thus, 300 μL of solid medium containing 0.5 % DMSO and 0–100 μM thaxtomin A was laid into each well of a clear-bottomed, clear-walled 96-well plate. *C. riethii* was scraped from an agar plate into liquid medium and the colonies sheared by pipetting. Cell density was adjusted to 2.5×10^6^ thalli mL^−1^ before pipetting 5 μL onto each solidified well. Chlorophyll absorbance was then measured daily as for *M. viride*.

### Light microscopy of plant roots

*N. benthamiana* seedlings were grown vertically on solid ½ MS medium until nine days after germination before transplanting to liquid ½ MS medium lacking sucrose and supplemented with 0.5 % DMSO ± 10 µM thaxtomin A. Rice seedlings were grown on solid ½ MS medium in plastic pots for five days after germination before transplantation.

Brightfield microscopy was performed with the EVOS M7000 imaging system at 10X magnification. Seedlings were directly mounted onto the slide in liquid ½ MS medium, over which a cover slip was placed. Multiple panels were acquired for each root before stitching using the Image Stitching plugin^60^ in FIJI^61^. For the time course video, a single live seedling was submerged in liquid ½ MS medium in a small coverglass-bottom petri dish which was placed into a vessel holder stage.

For calcofluor white staining, whole seedlings were stained for 30 min in 1 g L^−1^ calcofluor white M2R / 0.5 g L^−1^ Evans blue stain (#18909; Sigma-Aldrich) before direct mounting without washing. Root tips were then imaged at 10X magnification using a Zeiss LSM 880 confocal microscope using a 405-nm excitation laser and an emission range of 410–524 nm, collecting a Z-stack that was then summed using the Z project tool in FIJI.

Elongation zone cells within 150–250 μm of the transition zone with well-resolved cell wall features were then manually selected using the polygon tool in FIJI. Width, length, and aspect ratio were calculated using the in-built measurement tool. Statistical differences were assessed by one-way ANOVA in Rstudio version 2024.09.1.

### Leaf necrosis assay

Leaves of seven-week-old *N. benthamiana* plants were sprayed with a solution of 0.5 % DMSO and 0.5 % Tween-20 ± 10 µM thaxtomin A using a glass atomiser until run-off occurred. Plastic bags were secured over each sprayed leaf to increase humidity. Three plants were used per treatment. After 24 hours, leaves representing various stages of maturity were photographed.

### Cellotriose release assay

Three flasks were prepared per treatment. For each flask, 25 mg *N. benthamiana* seeds were germinated in 25 mL liquid ½ MS medium supplemented with 2 % sucrose. Five days after germination, the seeds were washed twice with 25 mL ½ MS medium lacking sucrose before the addition of a further 25 mL ½ MS medium (no sucrose) supplemented with 0.5 % DMSO ± 10 μM thaxtomin A. After three days, the medium was carefully harvested (taking care to remove any seeds) before centrifugation at 200,000×*g* for 40 min to remove debris. Control samples were also prepared consisting of ½ MS medium supplemented with 0, 10, 30, or 100 nM cellotriose or 0.5 % DMSO + 10 μM thaxtomin A.

The supernatants and control samples then underwent reverse phase chromatography as previously described^62^. For each sample, a HyperSep™ Hypercarb™ SPE Cartridge (25 mg bed volume; ThermoFisher Scientific) was treated with 1 M NaOH (0.5 mL) followed by water (1 mL), 30 % acetic acid (0.5 mL), and water (0.5 mL), followed by washing with 50 % acetonitrile supplemented with 0.1 % trifluoroacetic acid (solution A; 0.5 mL) and finally equilibration with 5 % acetonitrile supplemented with 0.1 % trifluoroacetic acid (solution B; 0.5 mL). Each sample was then passed through the column followed by washes with water (0.5 mL) and solution B (2 × 0.25 mL) and elution with solution A (4 × 0.2 mL). For PACE experiments (see below), 400 μL of the eluate was then dried by centrifugal evaporation before resuspension in 5 μL labelling master mix: 50 % DMSO, 7.5 % acetic acid, 50 mM 8-aminonaphthalene-1,3,6-trisulfonic acid (ANTS; ThermoFisher Scientific), and 50 mM 2-picoline borane and overnight incubation at 37 °C. After a second drying step, the samples were resuspended in 20 μL 6 M urea.

For β-glucosidase assays, two 100-μL aliquots of each eluate were dried by centrifugal evaporation before resuspension of each in 50 μL 100 mM ammonium acetate, pH 5.0 ± 0.5 μL *Aspergillus* sp. GH3-family β-glucosidase (Megazyme, Bray, Ireland). Both samples were then incubated at 37 °C overnight before drying, overnight incubation in 2.5 μL ANTS labelling master mix, drying again, and finally resuspension in 5 μL 6 M urea.

PACE was carried out as described previously^63^. To prepare standards, 2 μL of each 10 mM cello-oligosaccharide (Megazyme, Bray, Ireland) stock was combined before drying in a single tube and labelling with 5 μL labelling master mix as described above and resuspending in 100 μL 6 M urea. Samples in urea (2.5 μL each) were loaded onto a 240=×=180=×=0.75=mm polyacrylamide gel (29:1; acrylamide/bis) containing 0.1 M Tris-borate buffer, pH 8.2 (stacking gel: 10 % acrylamide; resolving gel: 20 % acrylamide) cast using the Hoefer SE660 electrophoresis system (Hoefer, Bridgewater, MA, USA) prior to electrophoresis in the same Tris-borate buffer at 200 V for 30 min followed by 1,000 V for 2 h. Gel fluorescence was then imaged using a Syngene G:Box Chemi XX6 imaging system equipped with a 365-nm wavelength UV transilluminator and short pass filter (516–600 nm).

### Ca^2+^ signalling assay

Eightlzldaylzlold seedlings grown in 96lzlwell plates were loaded overnight with 10=µM coelenterazine to allow aequorin reconstitution. Cytoplasmic Ca²⁺ influxes were recorded as Relative Light Units (RLUs) using a Varioskan LUX Reader luminometer (Thermo Scientific) following treatment with 10=µM thaxtomin A (in 5% DMSO or 1% methanol) or 10=µM cellotriose. Total Ca²⁺ discharge values were used to normalise Ca²⁺ bursts for each seedling.

### Insect cell expression and protein purification

Expression and purification of hybrid aspen and soy CesA proteins was carried out as described previously^4,29^. Baculovirus constructs (N-terminally His_10_-tagged *Ptt*CesA8 and mutants, *Gm*CesA1, *Hv*CslF6 and mutant, as well as N-terminally TwinStrep-tagged *Gm*CesA3 and *At*CslD3) were cloned into the pACEBac1 vector. Mutagenesis was carried out using the ‘QuickChange’ approach (mutagenic PCR). Constructs were transformed into DH10MultiBac cells and baculovirus was produced according to the NIH Joint Centre for Innovative Membrane Protein Technologies (JCIMPT) protocol (https://commonfund.nih.gov/sites/default/files/JCIMPT_LargeScalePurification.pdf). Sf9 cultures were grown in ESF921 medium at 27 °C with 145 rpm shaking. Expression was carried out in five or six 400 mL culture volumes infected using 15 mL P2 virus each. Cell pellets were harvested after 3–4 days at approximately 70–85 % viability.

For purification, 15–20 g of cell pellet (or 40 g for *At*CslD3) was homogenised with twenty strokes of a Dounce homogeniser in 120 mL membrane resuspension buffer (MRB; 20 mM Tris-HCl, pH 7.5, 100 mM NaCl, 5 mM sodium phosphate, 5 mM sodium citrate, 1 mM TCEP) with an additional 1% lauryl maltose neopentyl glycol (LMNG, Anatrace), 0.2% cholesteryl hemisuccinate (CHS, Anatrace), and protease inhibitors: 0.4 mM AEBSF, 2 μM aprotinin, 30 μM pepstatin, 7.5 μM pepstatin, 40 μM bestatin, 35 μM E-64, and 4.5 mM benzamidine hydrochloride. Following incubation on a rocker at 4 °C for 1 h, the solubilised samples were centrifuged at 200,000×*g* for 45 min. The supernatant then underwent affinity chromatography.

For His-tagged constructs, imidazole was added to a final concentration of 40 mM before batch incubation with 5 mL HisPure Ni-NTA resin (ThermoFisher Scientific) for 1 h at 4 °C on a rocker. Meanwhile, glyco-diosgenin (GDN, Anatrace) was added to MRB to a final concentration of 0.02 % to make buffer A. The Ni-NTA resin was transferred to a gravity flow column (20 × 2.5 cm Kimble FLEX-COLUMN; DWK Life Sciences, Wertheim, Germany) and washed with sixteen column volumes (CVs) of buffer A supplemented with 40 mM imidazole, 10 CVs of buffer A supplemented with 1 M NaCl, and 5 CVs buffer A supplemented with 60 mM imidazole. The protein was then eluted in 5 CVs buffer A supplemented with 400 mM imidazole.

For TwinStrep-tagged constructs, the supernatant was passed five times through 5 mL Strep-Tactin Sepharose resin (IBA Lifesciences, Göttingen, Germany) in a gravity flow column. The resin was then washed with 16 CVs buffer A, 10 CVs of buffer A supplemented with 1 M NaCl, and another 5 CVs buffer A. The protein was eluted in 5 CVs buffer A supplemented with 5 mM desthiobiotin.

Eluted protein was concentrated using Amicon centrifugal concentrators (Merck, Darmstadt, Germany) with a 100 kDa cut-off before size exclusion chromatography in buffer A using a Superose 6 Increase column. The relevant fractions were then concentrated to 16.6 μM for monomers / 5.5 μM for trimers.

BcsAB complex purified from *Cereibacter sphaeroides*, a gift of Dr Irek Górniak, University of Virginia, was expressed and purified as previously described^64^ except that the size exclusion buffer detergents were replaced with n-dodecyl β-maltoside (DDM) and CHS.

### Tritium incorporation assay

Tritium incorporation-based cellulose synthase activity assays were carried out as described previously^4,29,65^. For CesA / CslD reactions, 20-μL reactions were carried out in buffer A with 4.4 μM monomer / 1.5 μM trimer, 5 mM untritiated UDP-glucose, 0.34 μM ^3^H UDP-glucose (Perkin Elmer), and 20 mM EDTA or MgCl_2_ (in addition to inhibitor, prepared as described below). For BcsAB reactions, 20 μL reactions were carried out in 25 mM Tris-HCl, pH 7.5, 100 mM NaCl, 0.05 % DDM, 0.01 % CHS, and 0.01 % GDN with 10 μg mL^−1^ enzyme, 5 mM untritiated UDP-glucose, 0.34 μM ^3^H UDP-glucose (Perkin Elmer), 30 μM cyclic di-GMP, and 20 mM MgCl_2_. Reactions were incubated at 30 °C for 20 h (except when noted otherwise) before spotting onto filter paper in order to carry out descending paper chromatography in 65 % ethanol. Tritiated material retained at the origin was then quantified by liquid scintillation counting.

For inhibitor screens, reactions were additionally supplemented with 5 % DMSO and 0–0.5 mM inhibitor, diluted from 20 mM stocks in 100 % DMSO. For ‘DMSO-free’ reactions, 1.5 μL aliquots of 20 mM thaxtomin A in DMSO were dried at 60 °C in a centrifugal evaporator. The dried pellets were then resuspended in 30 μL buffer A (or 0.02 % GDN for BcsAB reactions) to make a 1 mM stock. This stock was diluted further in buffer A/0.02 % GDN to produce the necessary concentrations before combining with the reactions.

### Nano differential scanning fluorimetry

GDN-solubilised thaxtomin A was added at various concentrations to 400 nM *Ptt*CesA8 monomers. The mixtures were loaded into glass capillaries and a NanoTemper Tycho instrument (NanoTemper Technologies, Munich, Germany) was used to measure the thermal unfolding profile under default settings.

### Cryo-EM grid freezing and data collection

Purified protein was concentrated to 1.9 mg mL^−1^ (5.5 μM trimer). For thaxtomin-bound structures, aliquots of DMSO-solubilised thaxtomin were dried as above. Pellets were directly resuspended in concentrated protein by extensive pipetting to a final concentration of 4.5 mM before incubation on ice for 1 h. Any remaining insoluble material was briefly pelleted and removed. For structures bound to UDP- and UDP-Glc, 8 mM MgCl_2_ and 4 mM UDP or 4 mM UltraPure UDP-Glc (Promega, Madison, Wisconsin, USA) were added following the regular incubation with thaxtomin and incubated for an additional 30 min. Holey C-flat R1.2/1.3 300-mesh grids (Electron Microscopy Sciences) were glow-discharged in the presence of amylamine before the application of 2.5 μL protein at 4 °C and 100 % humidity and subsequent blotting and plunge-freezing using an FEI Vitrobot Mark IV.

Microscopy was carried out at the Molecular Electron Microscopy Core, University of Virginia. Grids were screened using a Thermo Fisher Glacios electron microscope prior to data collection on thin ice areas using a 300-kV FEI Titan Krios instrument. Data collection parameters are listed in Supplementary Table 1.

### Cryo-EM data processing

All datasets were processed in CryoSPARC^66^ v4.7.1. Processing flowcharts are available as **Supplementary Figs. 2, 5, 7, 8, and 13**. Movies were initially processed through patch motion correction and patch contrast transfer function (CTF) estimation followed by manual curation of micrographs to exclude exposures with poor CTF fit resolution (> 3.5–4 Å, depending on the dataset) and/or with defocus outside the accepted range (−2.0–−0.4 μm).

For the original, wild-type thaxtomin A-bound *Ptt*CesA8 structure (pixel size: 1.065 Å/pix), we used blob picking (200–280 Å diameter) to pick initial particles before subsequent 2D classification, generation of templates, and template picking. Picked particles were then binned fourfold before two rounds of 2D classification, *ab initio* reconstruction with three classes, and two rounds of 3D heterogeneous refinement using these classes as input. Because we found that 2D classification excludes particles with rare orientations at early stages, we then returned to the original template-picked particles and classified them directly with three rounds of 3D heterogeneous refinement, using the three classes generated in the first passthrough. After one round of 2D classification and removal of duplicate particles, the particles were re-extracted without binning and subjected to non-uniform refinement.^67^ For all non-uniform refinements, we applied optimisation of per-particle scale and no dynamic masking (by setting ‘dynamic mask start resolution’ to 1 Å). Importantly, no symmetry was applied. We then conducted further non-uniform refinements with CTF parameter optimisation following global CTF refinement and reference-based motion correction (including re-computation of dose weights), respectively. The result of the final non-uniform refinement constitutes the ‘consensus map’ of the *Ptt*CesA8 trimer.

After building initial structures into this consensus map, we realised that, although the individual protomers are identical, their tilt is somewhat variable within the trimeric assembly, inconsistent with C3 symmetry and preventing good alignment of the transmembrane region between different particles. Therefore, we implemented a symmetry expansion approach. Accordingly, the consensus map was aligned to the cyclic symmetry axis prior to C3 symmetry expansion. This created three copies of each particle, each with rotations of 0°, 120°, or 240° around the symmetry axis, respectively, thus superimposing each protomer on both of its neighbors. We then created a focus mask around one protomer using the ‘molmap’ command in UCSF Chimera^68^ version 1.17.3, which (after dilating and padding in CryoSPARC) was used to run local refinement on the expanded particles, producing a ‘focused map’ of one set of superimposed protomers.

Using the previous model^4^ of *Ptt*CesA8 (PDB: 6WLB) as a starting point, we then built and refined an atomic model of this protomer density using ISOLDE^69^ version 1.10.1. Thaxtomin A and C models and restraints were created using eLBOW^70^ in Phenix^71^ version 1.21.1-5286 under default settings. In the absence of true atomic resolution, we manually restrained coplanarity of the nitro group and indole ring. Furthermore, we modelled well-resolved water molecules within ∼7 Å of ligands of interest. Finally, in order to refine bond angles and lengths, the overall model underwent a limited real-space refinement^72^ in Phenix version 2.0-5885 using restraints provided by the ‘isolde write phenixRsrInput’ command in ISOLDE.

To produce the final trimeric structure, we combined three copies of this protomeric model after aligning them with the three protomeric densities of the original consensus map. This model was then used to accurately align and merge three copies of the focused map using the ‘combine focused maps’ tool in Phenix version 1.21.1-5286 to produce the final ‘composite map’.

In the other datasets (pixel size: 0.652 Å/pix), we followed an almost identical scheme, but bypassed unnecessary blob picking and 2D classification steps through the use of projection-based template picking. For the H832W mutant structure, we produced picking templates from the WT structure and repeated all steps from the 2D classification stage. For the remaining three datasets, we not only used the H832W structure to make picking templates, but recycled classes from heterogeneous refinement to begin directly with heterogeneous refinement of the template-picked particles. For these four datasets, local CTF refinement was also beneficial after global CTF refinement.

### Computational chemistry

The Gaussian 16^73^ set of programs was employed to carry out the quantum chemical calculations, applying the density functional theory (DFT) M06-2X functional^74^ in conjunction with the triple-ζ def2-TZVP basis set. Geometries were fully optimised, and characterised as true minima, by harmonic vibrational analysis which yielded all positive frequencies. The interaction energy is equal to the difference between the energy of the dyad and the sum of the energies of the two monomers in the geometry they adopt within the complex, then corrected for basis set superposition error by the standard counterpoise prescription^75^. The influence of a surrounding polarisable dielectric continuum made use of the polarisable continuum model (PCM) method incorporated into Gaussian.

### UDP-Glo assays

For UDP-Glo assays, thaxtomin A and thaxtomin C (sc-364220; SantaCruz Biotechnology, Inc., Dallas, TX, USA) dilution series were prepared in DMSO using a Hamilton syringe. Reactions (25 μL volume) containing 1 nM monomeric enzyme, 20 mM MgCl_2_, 5 % DMSO, and 0–1 μM thaxtomin A in buffer A were incubated at 30 °C for 2 h in a white, non-binding 96-well plate (Greiner Bio-One, Monroe, North Carolina, USA). Subsequently, 0–1 mM (for Michaelis-Menten) or 150 μM (for determination of inhibition constant) UltraPure UDP-glucose (Promega) was added. Following further incubation at 30 °C for 1 h, 25 μl UDP Detection Reagent (UDP-Glo kit; Promega) was added before a final incubation at room temperature. After 1 h, luminescence was detected using a GloMax Explorer plate reader (Promega).

### Tryptophan fluorescence quenching assay

For tryptophan fluorescence quenching assays, thaxtomin A dilution series were prepared in DMSO using a Hamilton syringe. Solutions (1 mL volume) containing 10 nM monomeric enzyme, 5 % DMSO, and 0–10 μM thaxtomin A were nutated at room temperature in Protein LoBind microcentrifuge tubes (Eppendorf, Hamburg, Germany). After 2 h, each solution was loaded in turn into the same quartz cuvette (scrupulously clean), and the tryptophan fluorescence emission spectrum (Ex/Em = 283/305–365 nm) was recorded using a FluoroLog-3 spectrophotometer. The blank spectrum for water (including the Raman scattering peak at 314 nm) was subtracted from each experimental spectrum. Total Trp fluorescence was measured as the sum of fluorescence emitted between 325 and 345 nm. Three independent technical replicates (separate days) were carried out for each data point.

### Calculation of Michaelis-Menten, inhibition, and binding constants

For Michaelis-Menten experiments, initial reaction velocity (UDP product accumulation in the first hour, confirmed to be in the linear phase) was plotted against substrate concentration. Michaelis-Menten parameters were fitted in GraphPad Prism version 6.0g.

For calculation of apparent inhibition constants, initial reaction velocity (*v*_i_) was plotted against total inhibitor concentration ([*I*]_T_). Using GraphPad Prism, the data were modelled according to the following equation^32^:

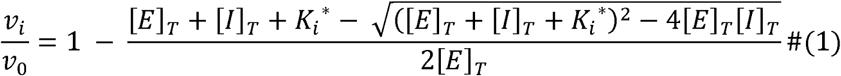

In these calculations, [*E*]_T_ was constrained at 1 nM, whereas *v*_0_ and 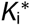 were fitted computationally.

To deduce binding constants, fluorescence was plotted against total inhibitor concentration, before modelling the data according to equation 1 (replacing 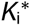 with dissociation constant, *K*_d_).

## Data availability

The following cryo-EM maps and models are available at the Electron Microscopy Data Bank and Protein Data Bank respectively: *Ptt*CesA8 bound to thaxtomin A (EMD-75083; PDB_000010DE), *Ptt*CesA8 bound to thaxtomin C (EMD-75084; PDB_000010DF), *Ptt*CesA8 bound to thaxtomin A and UDP-glucose (EMD-75085; PDB_000010DG), *Ptt*CesA8 bound to thaxtomin A and UDP (EMD-75086; PDB_000010DH), and *Ptt*CesA8[H832W] bound to thaxtomin A (EMD-75087; PDB_000010DI).

## Acknowledgements

We thank Dr Balasubramanyam Chittoor for initial exploratory findings; Dr Michael Purdy and Dr David Cooper (Molecular Electron Microscopy Core facility, University of Virginia) for electron microscopy support; Dr Mark Daniels (University of Virginia) for design, construction, and maintenance of essential equipment; Prof. Michael Timko and Dr Hai Liu (University of Virginia), Prof. Nam-Chon Paek (Seoul National University), and the MCC-NIES (Japan) for provision of plant and algal material; Mr Christopher Claussen and the University of Virginia Department of Biology greenhouse team for horticultural support; Dr Ireneusz Gόrniak (University of Virginia) for purification of bacterial cellulose synthase; and Prof. Zygmunt Derewenda (University of Virginia), Dr Henry Temple (Sainsbury Laboratory Cambridge University), and Dr Valentin Delobel (University of Virginia) for constructive scientific discussions.

This work was supported by a grant to S.S. by the US National Science Foundation (1954310). L.W., C.L., and P.P. were supported by the Howard Hughes Medical Institute, of which J.Z. is an investigator. Y.W. and R.H. were supported by NIH grant R35GM144130, awarded to J.Z.. MAT is supported by grants PID2021-126006OB-I00 and PID2024-159175OB-I00 from the Spanish Ministry of Science and Innovation. Transmission electron micrographs were recorded at the University of Virginia Molecular Electron Microscopy Core facility (RRID: SCR_019031), which is supported in part by the School of Medicine. In addition, the Titan Krios (SIG S10-RR025067) and K3/GIF (U24-GM116790) were purchased in part or in full with the designated NIH grants.

This article is subject to HHMI’s Immediate Access to Research policy, which requires that this article be made publicly available as initial and revised preprints deposited on a designated preprint server under a CC BY 4.0 license.

## Author contributions

Conceptualisation: LW, JZ; Formal analysis: LW, CL, SS; Investigation: LW, CL, MAT, RH; Methodology: LW, CL, MAT, SS, RH, ZD, JZ; Resources: LW, CL, MAT, YW, PP, ZD; Supervision: JZ; Writing – original draft: LW; Writing – review & editing: all

## Figure legends

**Extended Data Fig. 1.**
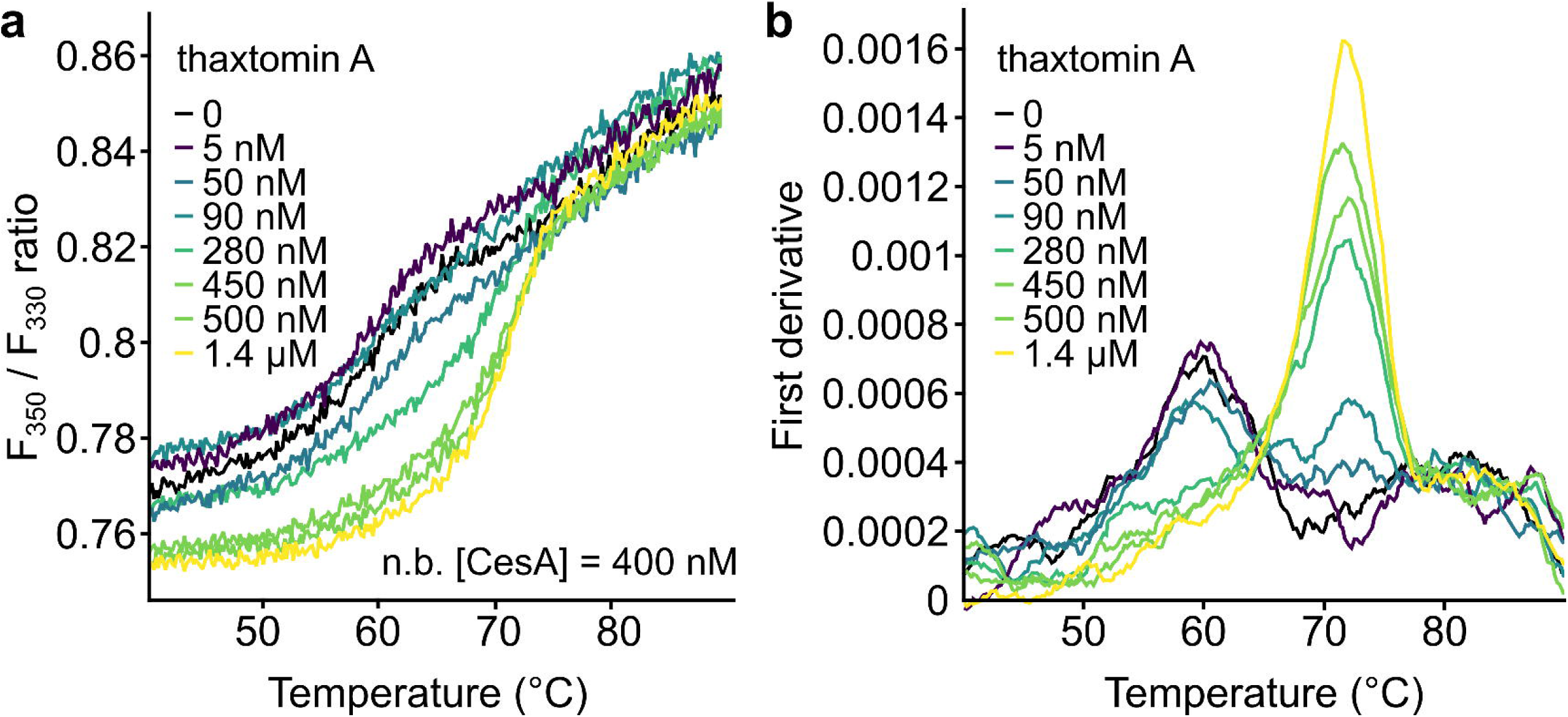
Detection of thaxtomin binding by nano-differential scanning fluorimetry (nanoDSF) Thermal unfolding profiles of monomeric *PttCesA8* were monitored according to the ratio of tryptophan fluorescence emission at 350 and 330 nm at varying concentrations of thaxtomin A. Protein concentration was maintained at 400 nM. **a** Unfolding profiles. **b** First derivative plots; showing fastest points of unfolding.

**Extended Data Fig. 2.**
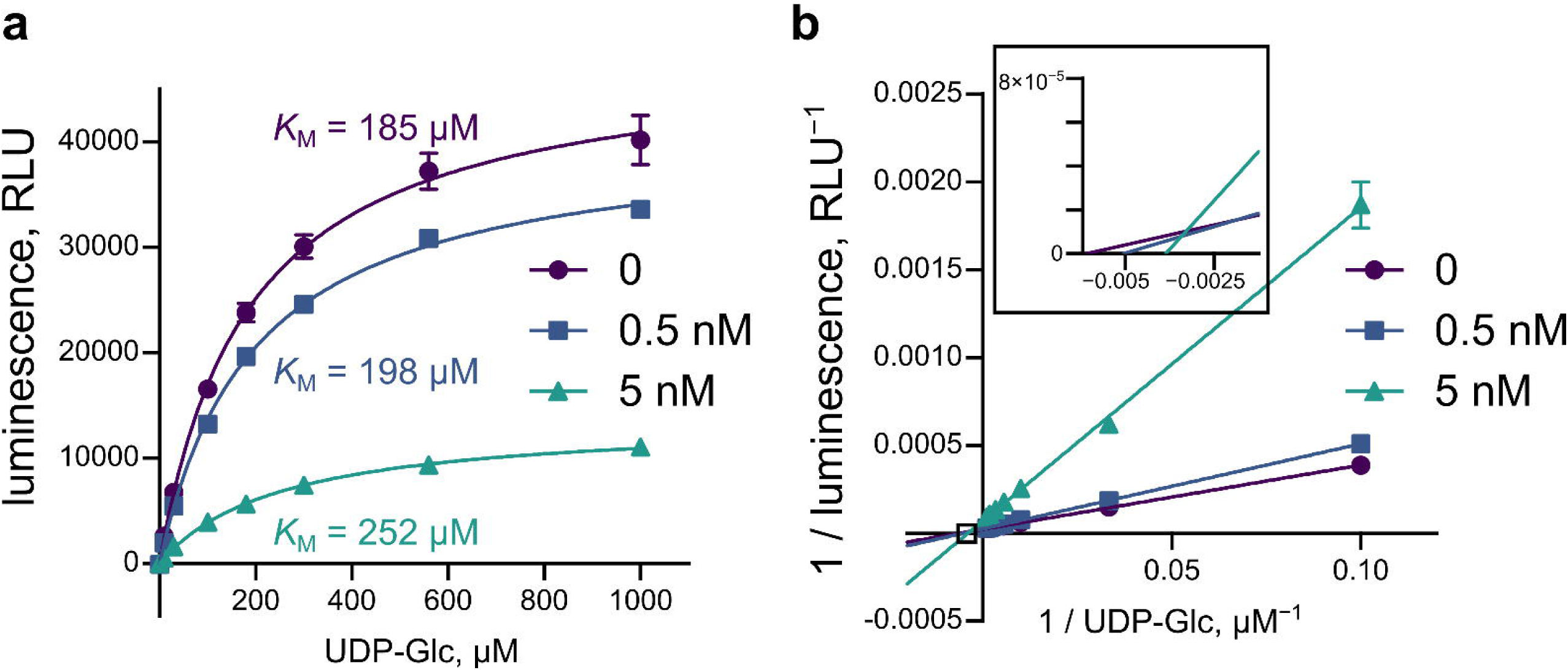
Effect of thaxtomin A on PttCesA8’s responsiveness to UDP-glucose. Inhibition of PttCesA8 activity was observed using the UDP-Glo assay. Enzyme (1 nM) was pre-incubated with 5 % DMSO ± thaxtomin A for 2 h before incubation with UDP-glucose substrate for 1 h at 30 °C. Free UDP is converted into luminescence. a Michaelis-Menten data recorded at different concentrations of inhibitor. b Lineweaver-Burke transformation of data shown in panel A.

**Extended Data Fig. 3.**
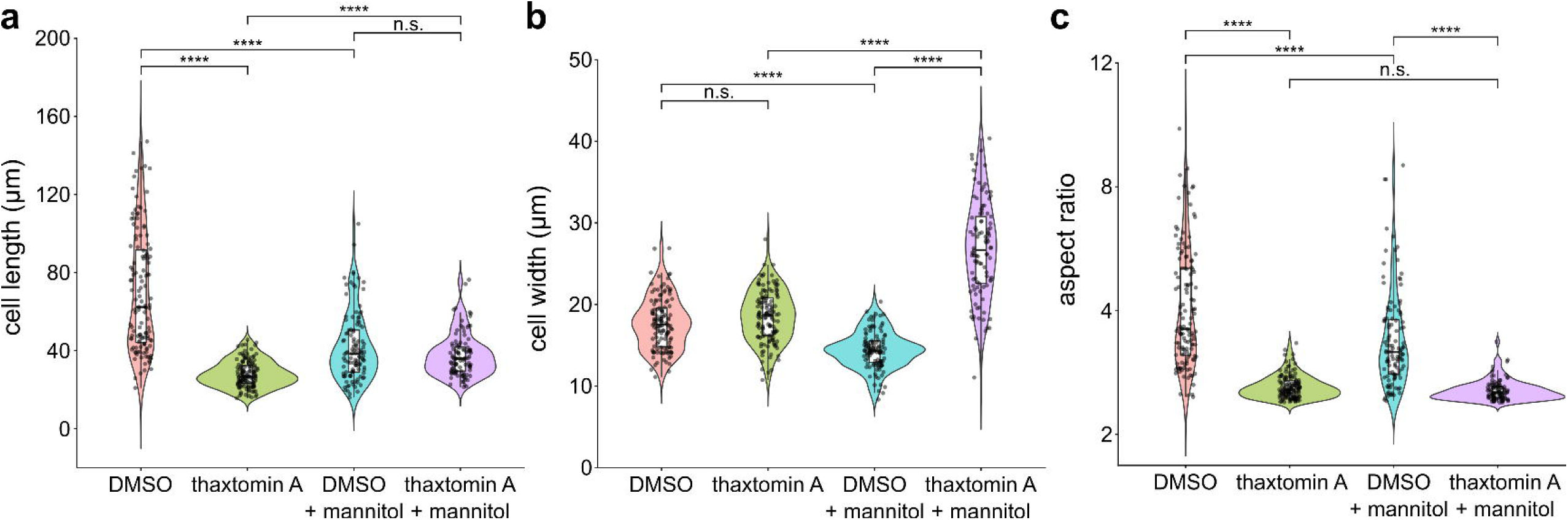
Cell measurements of N. benthamiana root expansion zone epidermal cells after treatment with or without thaxtomin. **A** Nine-day-old N. benthamiana seedlings were transplanted to liquid medium containing 0.5 % DMSO with or without 10 µM thaxtomin A and/or 0.3 M mannitol. After 24 hours, cell walls were stained with calcofluor white prior to confocal microscopy. The dimensions of epidermal expansion zone cells (23–31 cells per plant, five (no mannitol) or four (mannitol) plants per treatment; see Methods) were measured using FIJI. a Cell lengths (longest dimension). b Cell widths (shortest dimension). c Length:width aspect ratio. Statistics: one-way ANOVA; n.s. no significant difference; **** p < 0.0001.

