## Supplementary Information for "A molecular description of plant cellulose biosynthesis inhibition"

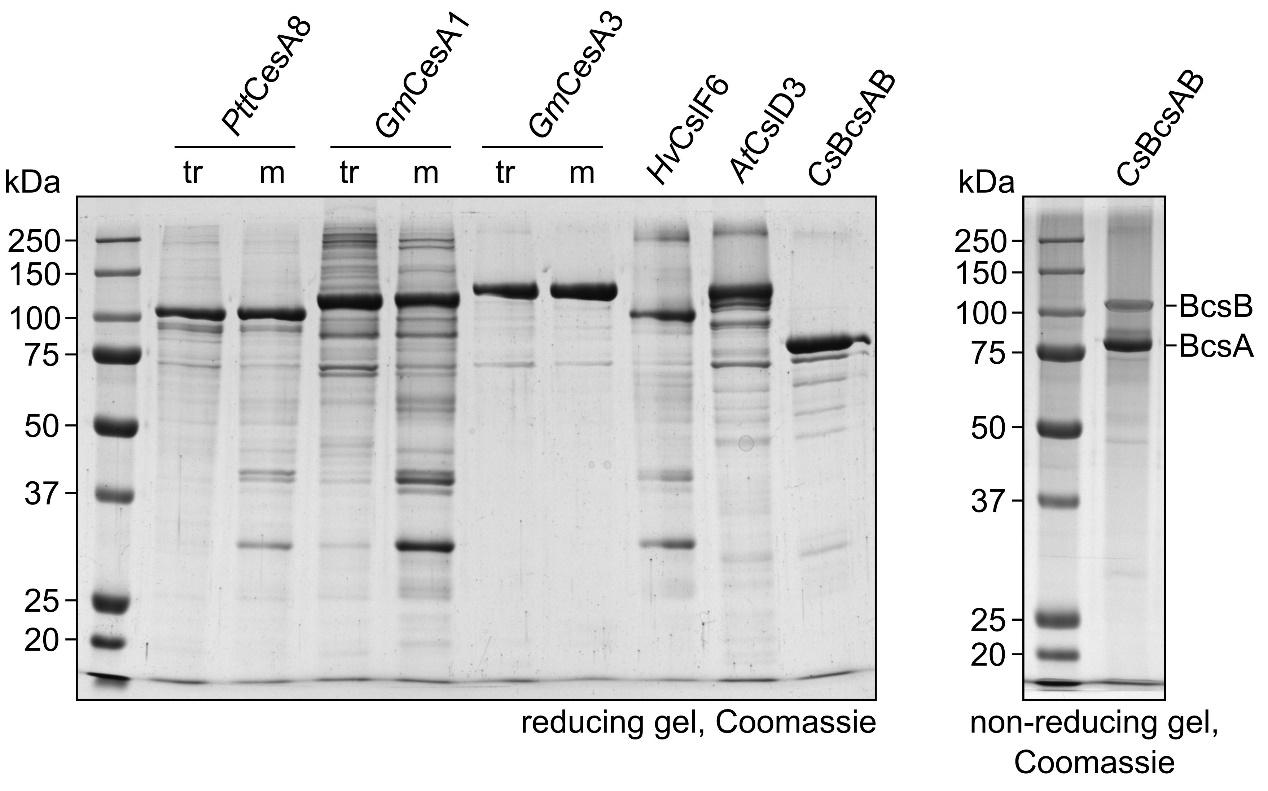

**Supplementary Fig. 1. SDS-PAGE analysis of purified cellulose synthase(-like) enzymes.**

Trimer: ‘tr’; monomer: ‘m’. Expected molecular weights: His_12_–*Ptt*CesA8: 112.3 kDa; His_12_–GmCesA1: 123.9 kDa; TwinStrep–GmCesA3: 123.6 kDa; His_12_–*Hv*CslF6: 106.8 kDa; TwinStrep–*At*CslD3: 131.5 kDa; *Cs*BcsA–(His_6_)_2_: 89.9 kDa; *Cs*BcsB: 74.7 kDa. Note that *Cs*BcsA–(His_6_)_2_ and *Cs*BcsB co-migrate under reducing conditions.

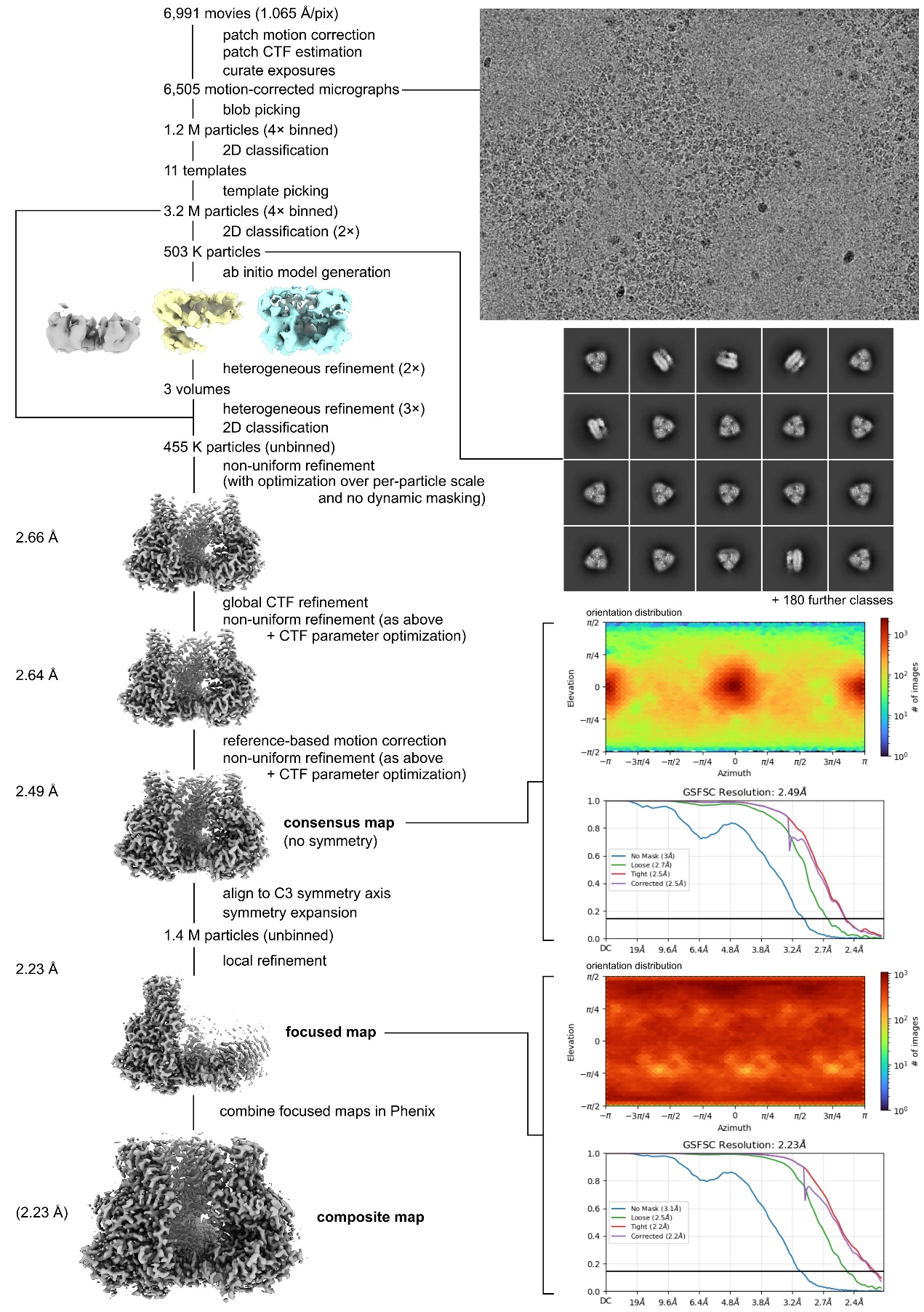

**Supplementary Fig. 2. CryoSPARC processing strategy to determine the structure of *Ptt*CesA8 bound to thaxtomin A**

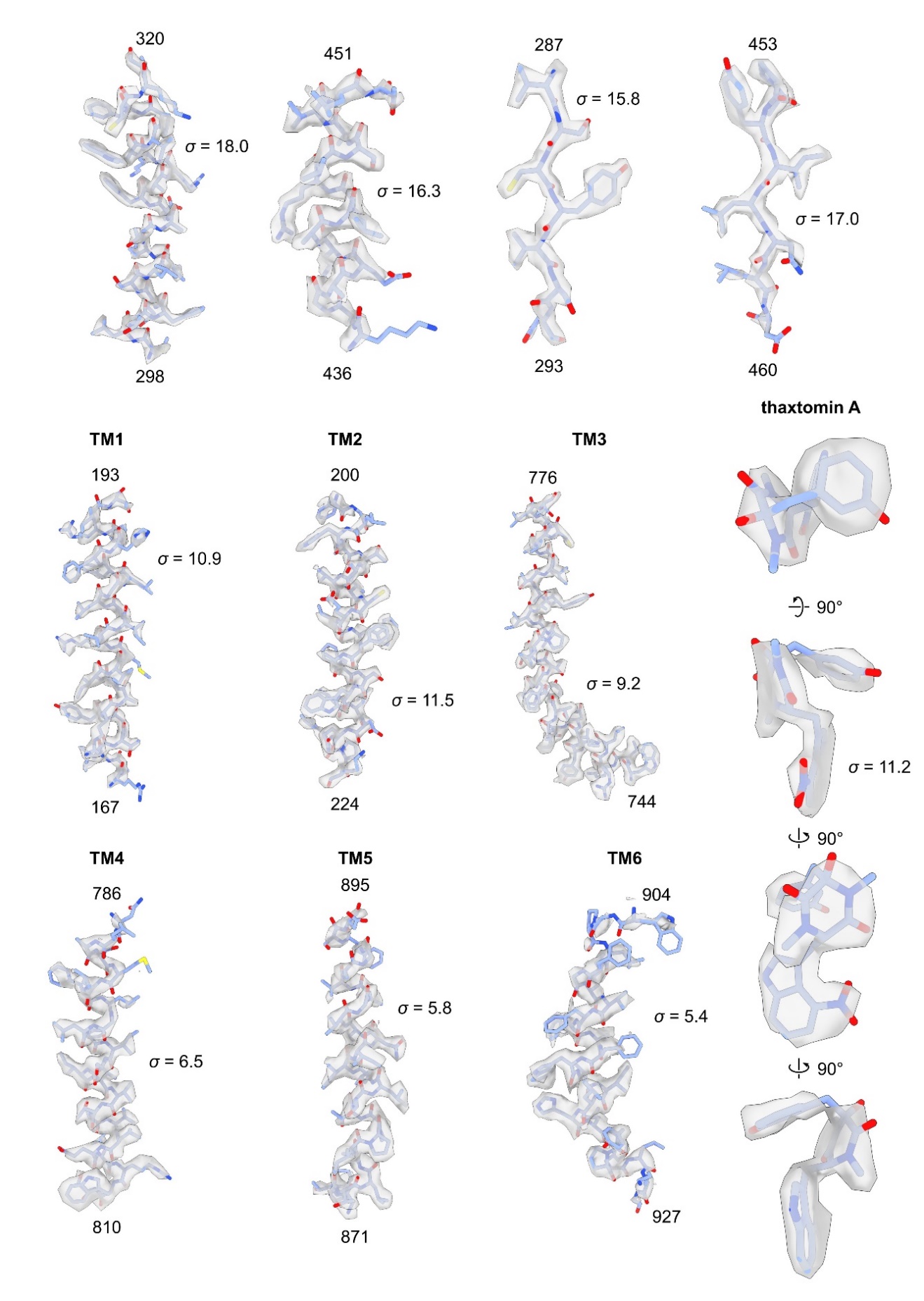

**Supplementary Fig. 3. Local cryo-EM density for selected regions of *Ptt*CesA8 + thaxtomin A map**

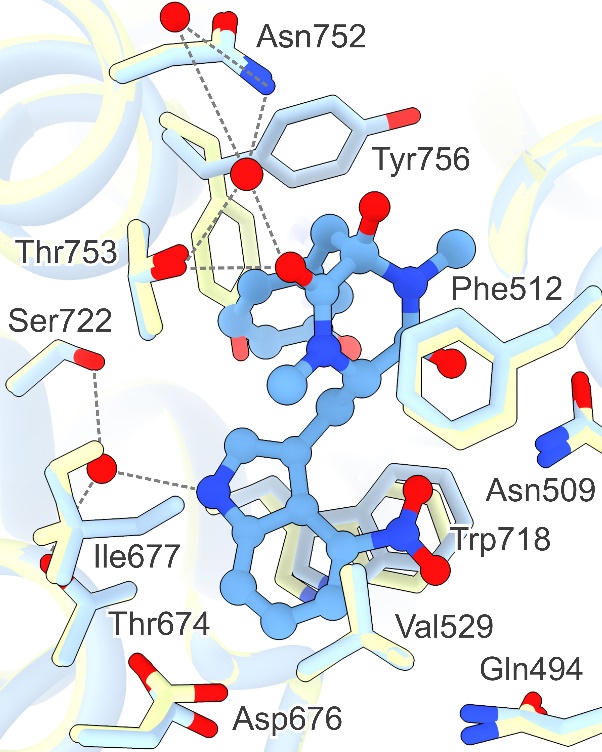

**Supplementary Fig. 4. Water coordination in the thaxtomin A binding site**

Structure of *Ptt*CesA8 bound to thaxtomin A (blue), aligned to structure of apo *Ptt*CesA8 (lime green). Waters shown as red spheres. Interactions involving bound water molecules are highlighted as dashed lines.

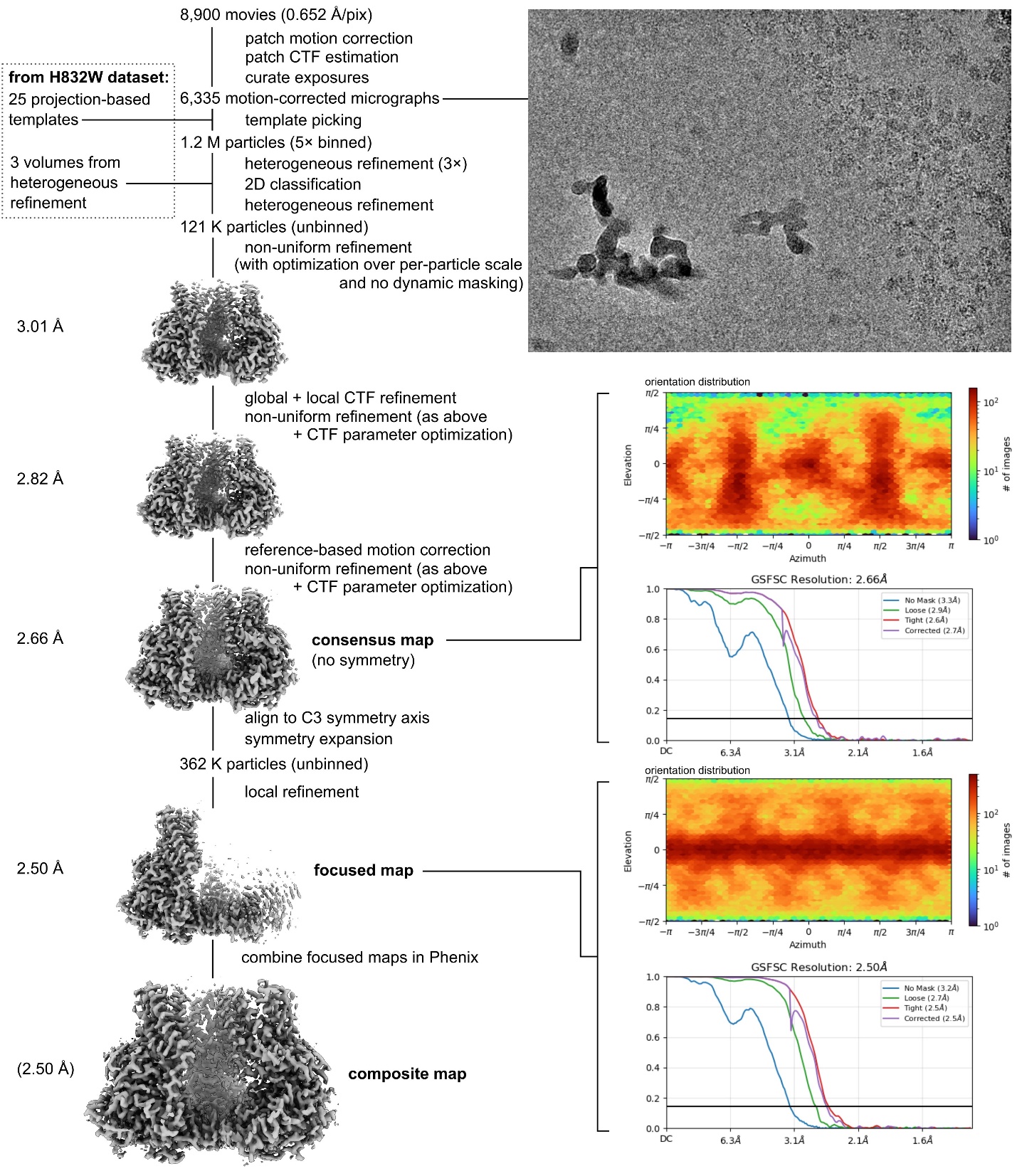

**Supplementary Fig. 5. CryoSPARC processing strategy to determine the structure of *Ptt*CesA8 bound to thaxtomin C**

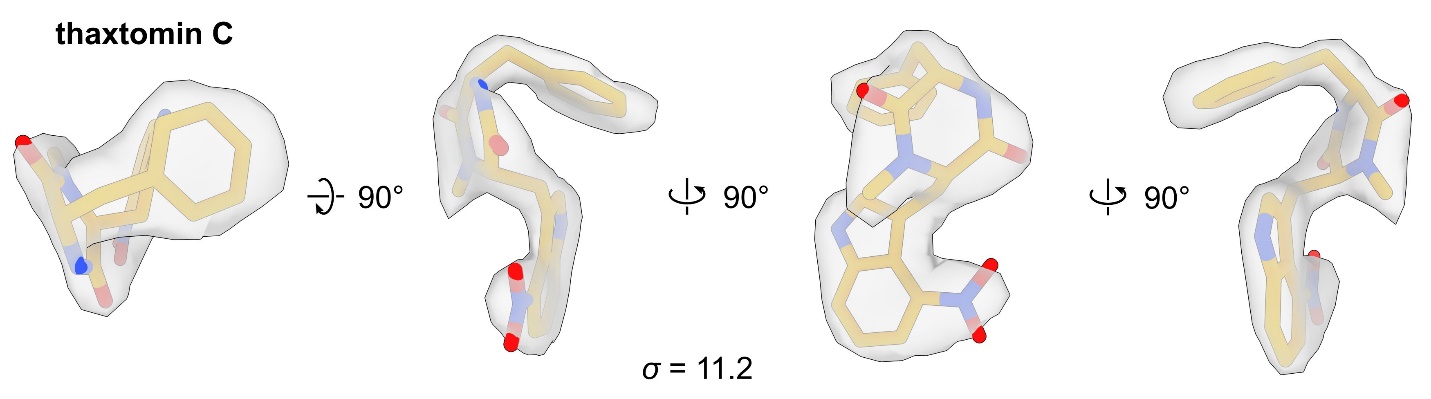

**Supplementary Fig. 6. Local cryo-EM density for thaxtomin C**

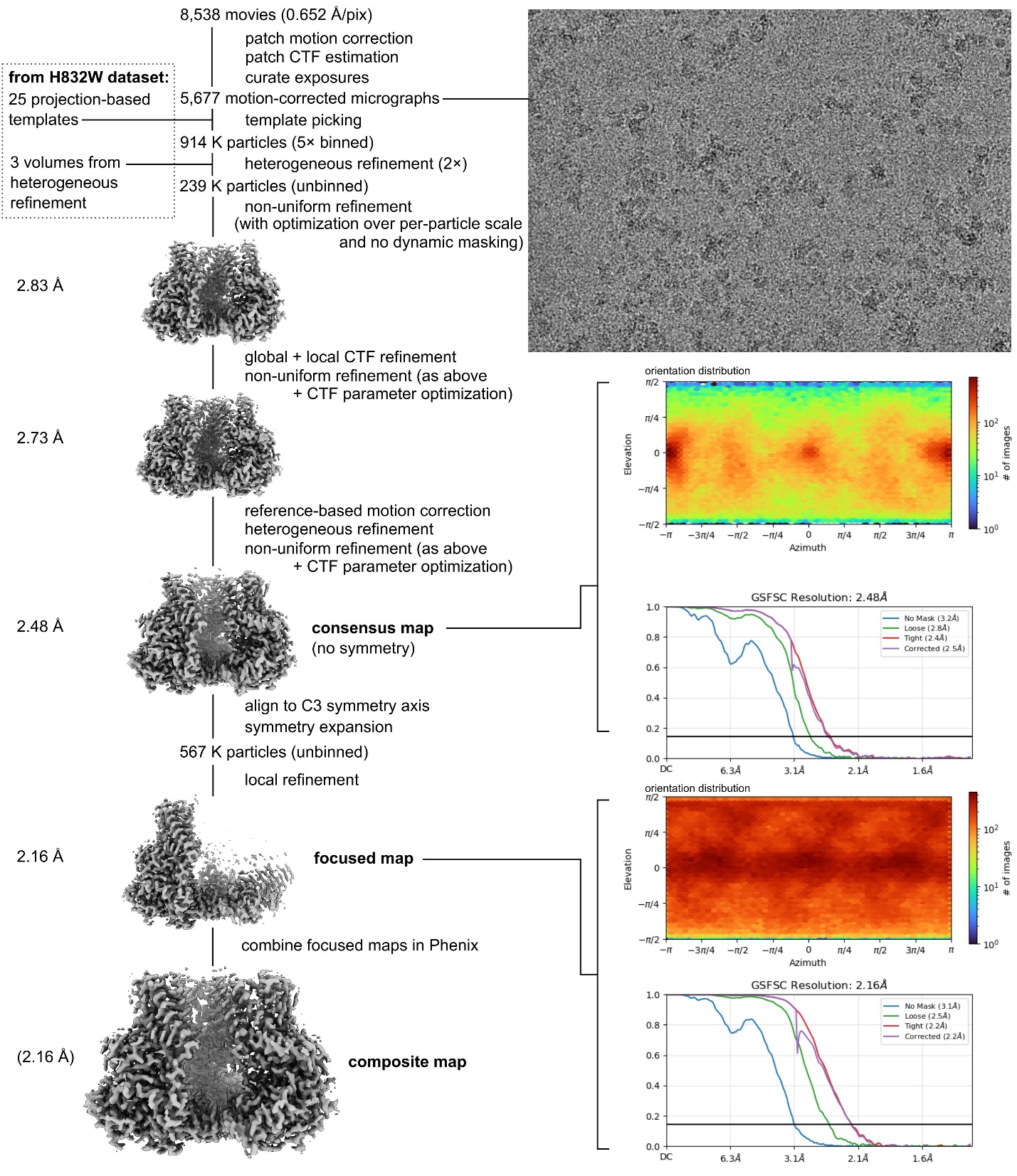

**Supplementary Fig. 7. CryoSPARC processing strategy to determine the structure of *Ptt*CesA8 bound to thaxtomin A and UDP-glucose**

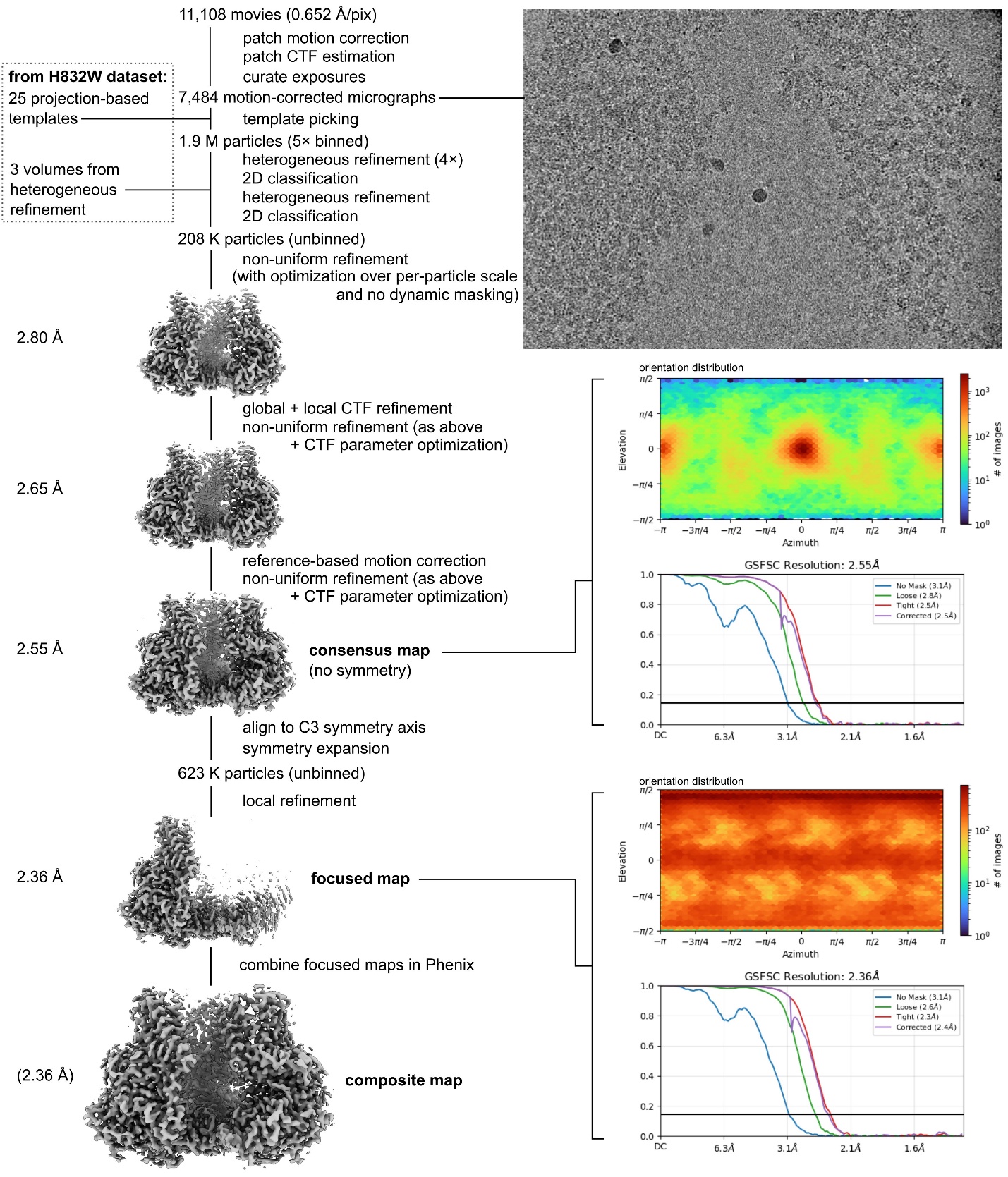

**Supplementary Fig. 8. CryoSPARC processing strategy to determine the structure of *Ptt*CesA8 bound to thaxtomin A and UDP**

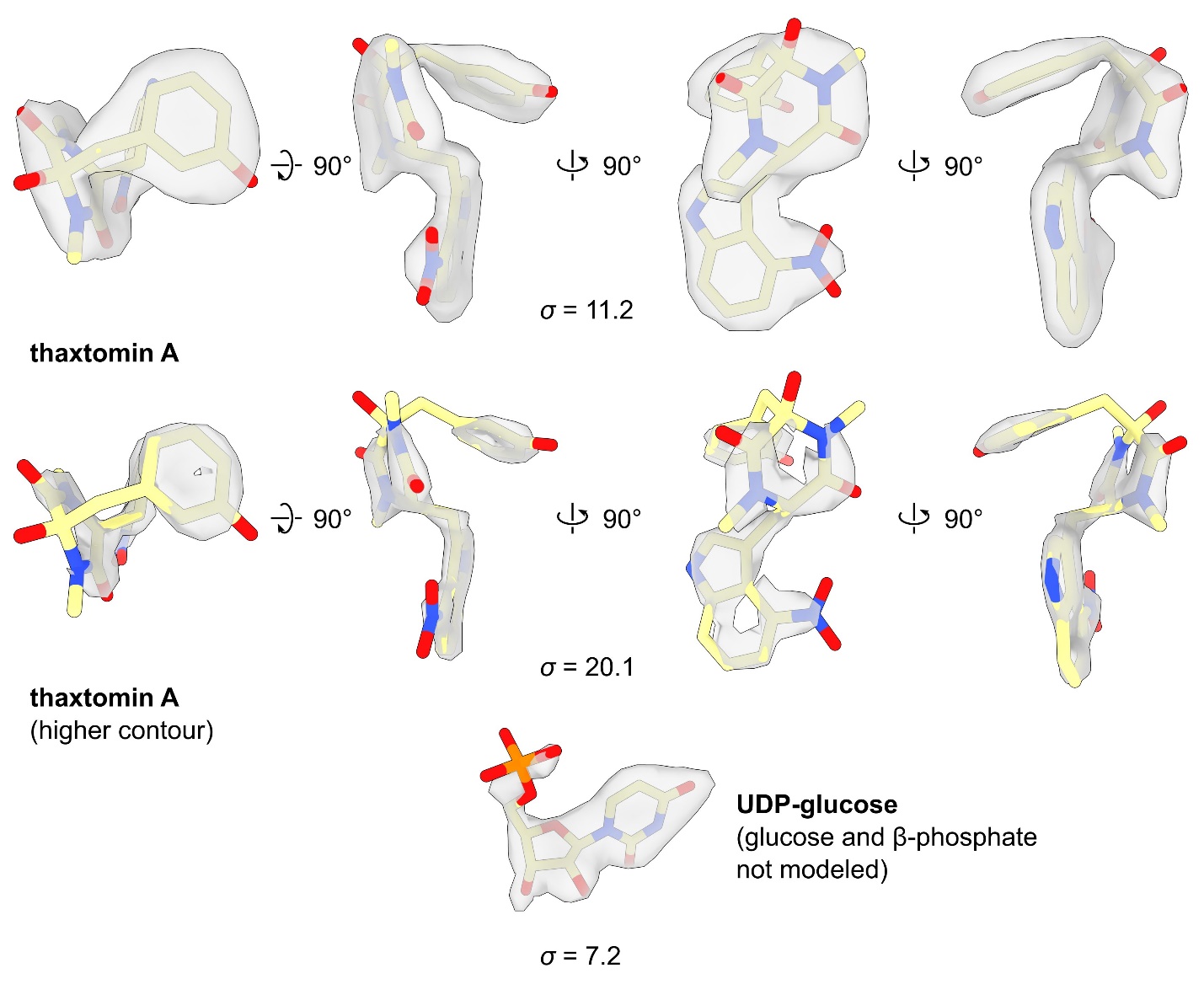

**Supplementary Fig. 9. Local cryo-EM density for ligands observed in *Ptt*CesA8 + thaxtomin A + UDP-glucose structure, highlighting high resolution details visible for thaxtomin A**

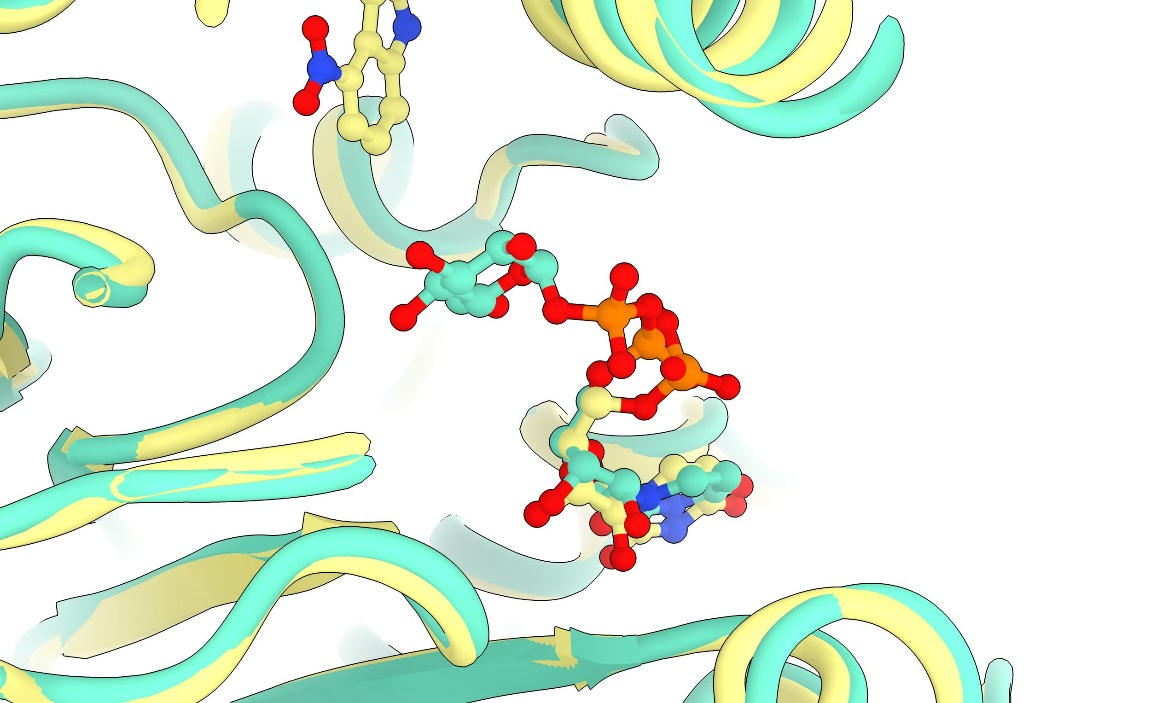

**Supplementary Fig. 10. Structural alignment of *Ptt*CesA8 bound to UDP-glucose (green; PDB: 8G2J) with *Ptt*CesA8 bound to UDP-glucose and thaxtomin A (yellow; this work).**

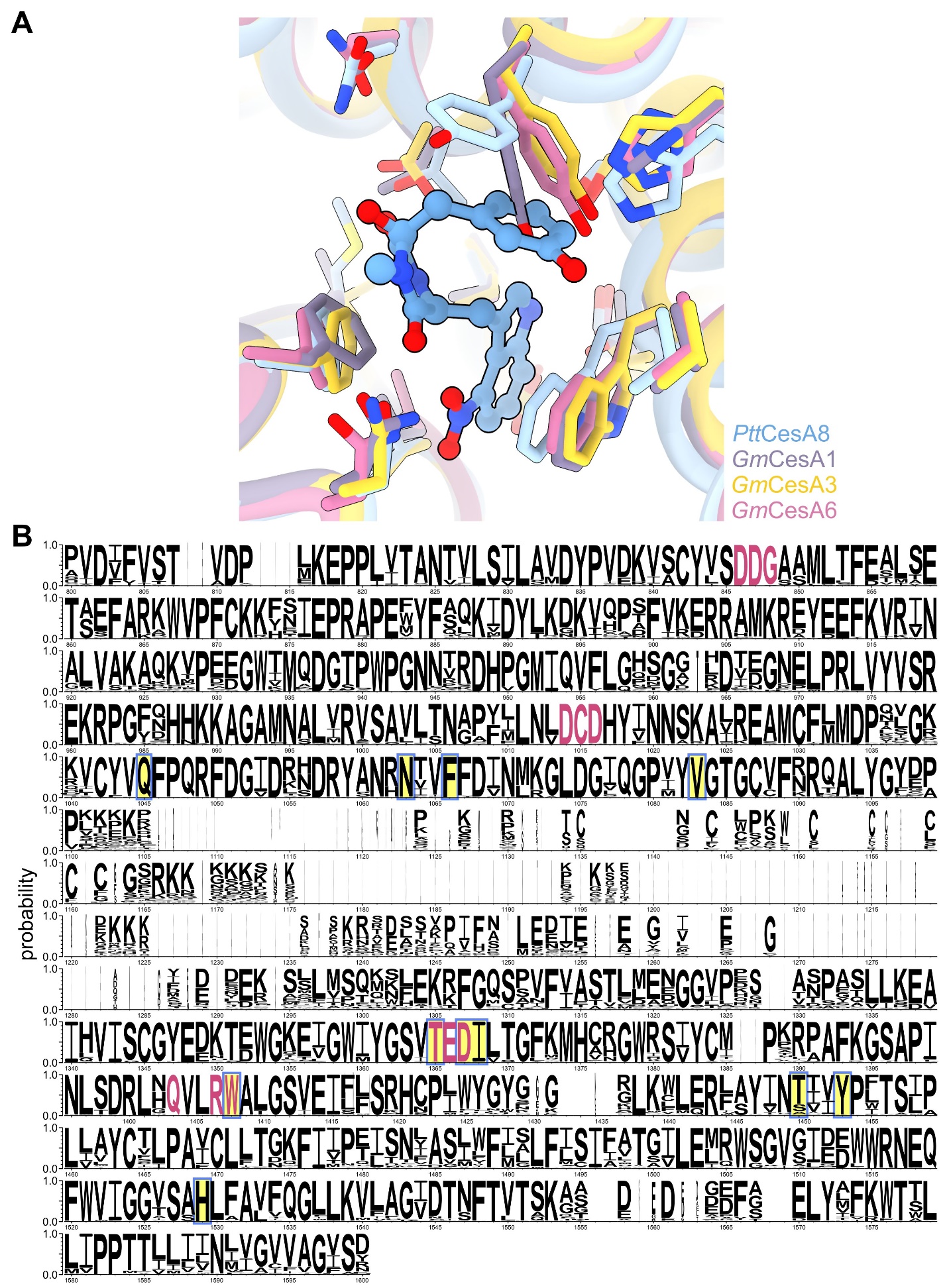

**Supplementary Fig. 11. Conservation of the thaxtomin binding site**

**A** Structural alignment of experimental structures of *Ptt*CesA8 bound to thaxtomin A (blue), *Gm*CesA1 (8VHZ; purple), *Gm*CesA3 (8VHT; yellow), and *Gm*CesA1 (8VI0; pink). **B** Sequence logo showing residue conservation amongst 64 CesA sequences from a diverse range of plant species (Arabidopsis, poplar, rice, *Amborella trichopoda*, giant redwood, *Physcomitrium patens*, and *Chara braunii*), showing positions 800–1,600 (corresponding to residues 252–895 of *Ptt*CesA8) of the alignment (1,693 columns in total). Sequences were obtained from Dicots PLAZA 5.0 (https://vandepoelelab.be/plaza/versions/plaza_v5_dicots/) and aligned with Muscle5. After removal of sequences with long indels and re-alignment, the sequence logo was created with WebLogo 3 (https://weblogo.threeplusone.com/). Residues directly adjacent to thaxtomin A in the *Ptt*CesA8 structure are highlighted in yellow, and previously characterized sequence motifs are coloured in pink.

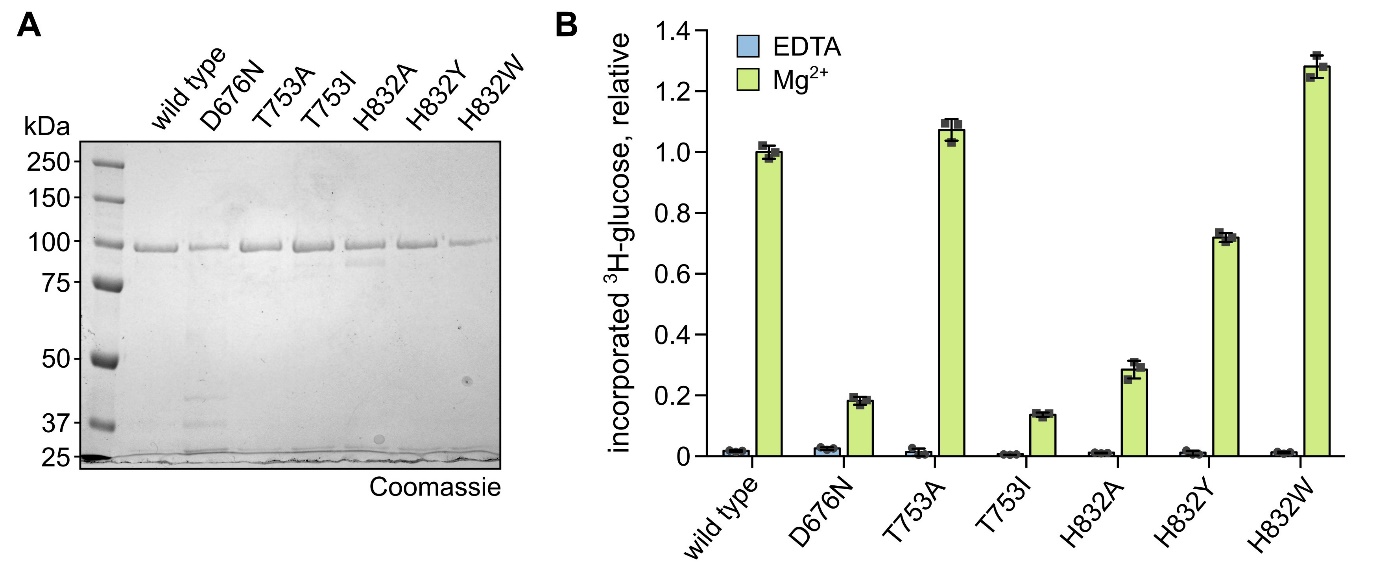

**Supplementary Fig. 12. Purification and activity of *Ptt*CesA8 mutants**

**A** Purity of the various *Ptt*CesA8 mutants as assessed by SDS-PAGE, stained with Coomassie. Expected molecular weight of His_12_–*Ptt*CesA8: 112.3 kDa. **B** Activity of the mutants (90 min reaction time) as characterized by tritium incorporation assay, normalized to wild-type reaction. EDTA replaced Mg^2+^ in negative control reactions. Three technical replicates were conducted.

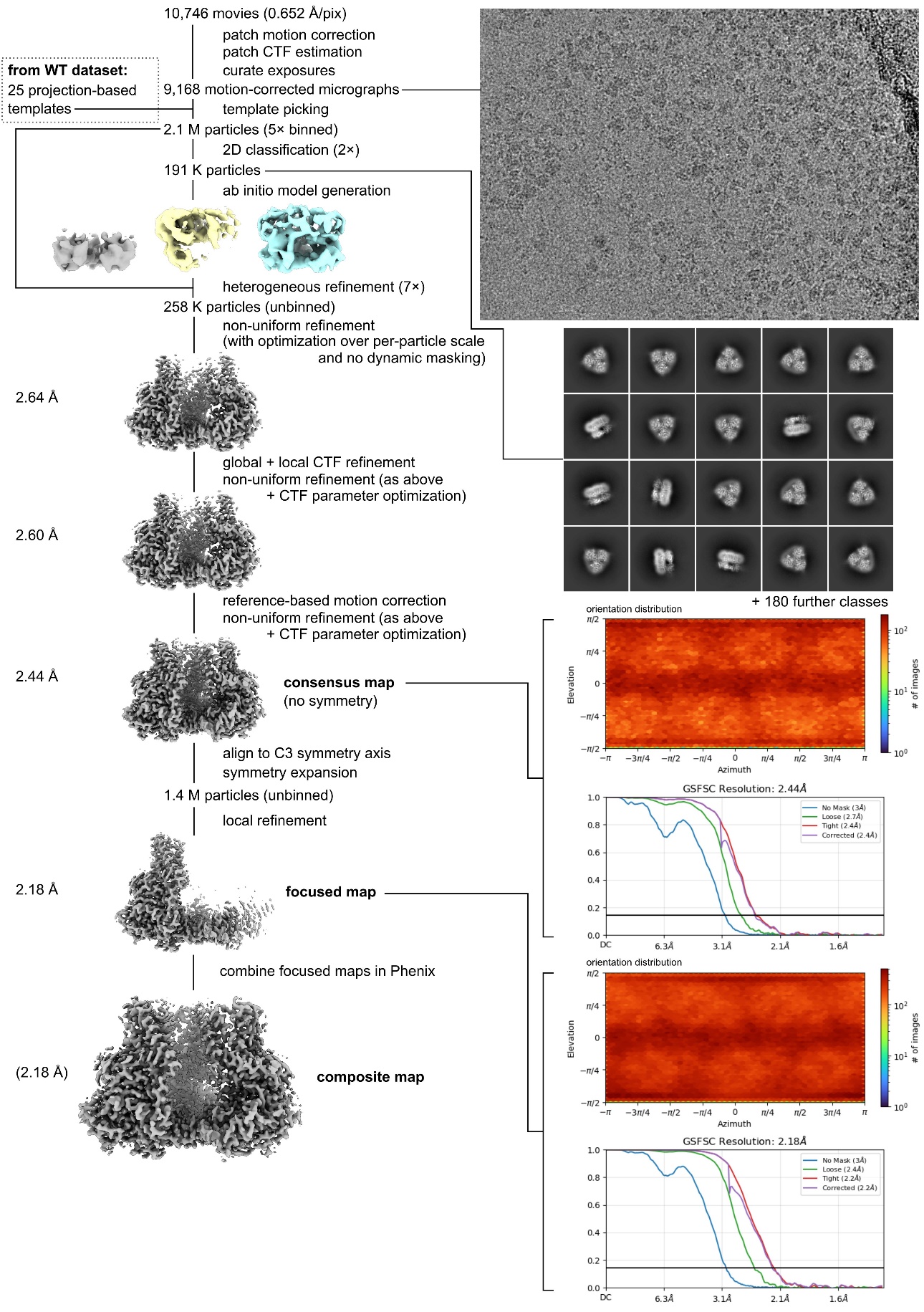

**Supplementary Fig. 13. CryoSPARC processing strategy to determine the structure of *Ptt*CesA8 H832W mutant bound to thaxtomin A**

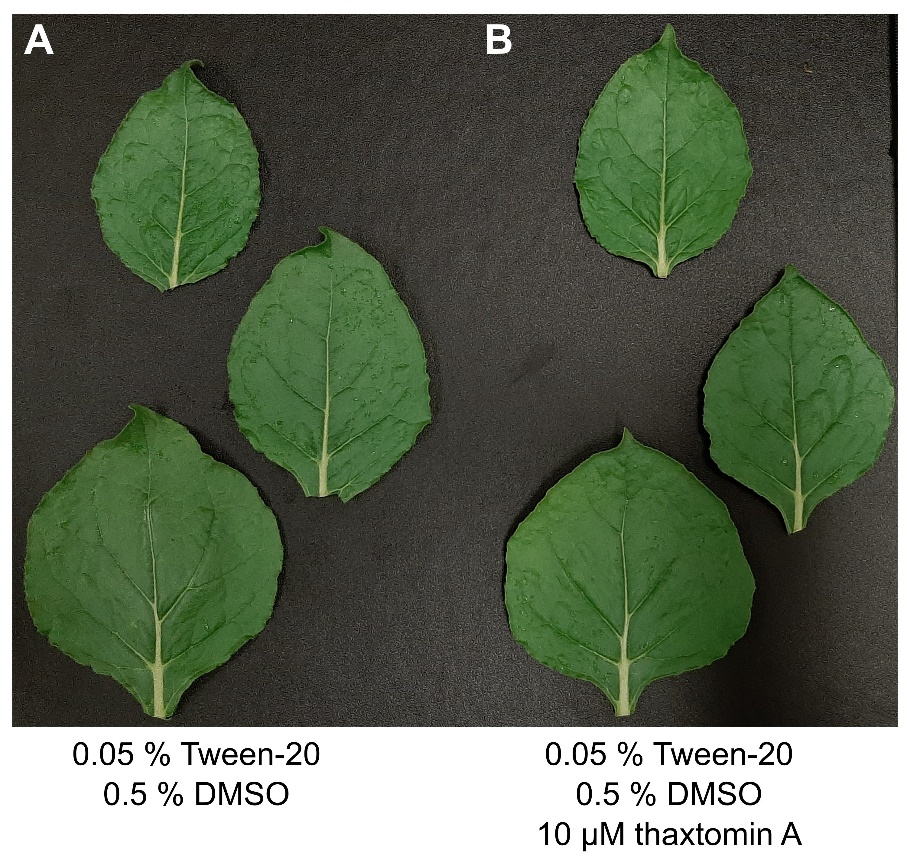

**Supplementary Fig. 14. Responses of *Nicotiana benthamiana* leaves to thaxtomin A exposure.**

Seven-week-old *N. benthamiana* plants were sprayed with a solution of Tween-20 and DMSO with or without 10 μM thaxtomin A until run-off occurred. Representative leaves from three plants were photographed after 24 hours.

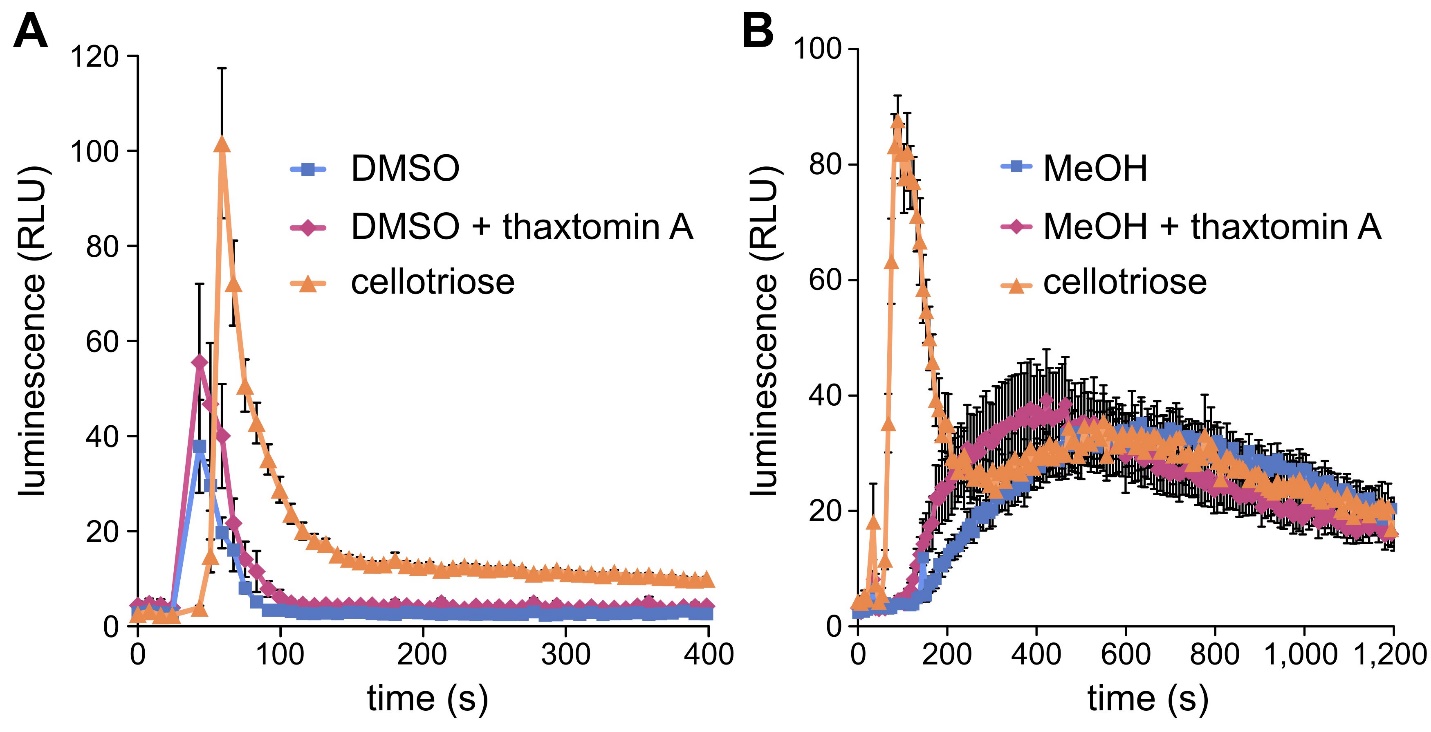

**Supplementary Fig. 15. Thaxtomin A does not convincingly elicit a global cytoplasmic Ca^2+^ burst in Arabidopsis seedlings compared with solvent alone**

Nine-day-old liquid-grown Arabidopsis seedlings expressing the apoaequorin Ca^2+^ reporter (Col-0^AEQ^) were treated with various chemical stimuli. Immediate changes in cytoplasmic Ca^2+^ concentration were monitored as luminescence, normalized to the total Ca^2+^ discharge for each seedling. The damage-associated molecular pattern cellotriose served as a positive control for the Ca^2+^ burst response. **A** Plants were treated with 0.5 % DMSO, 0.5 % DMSO + 10 µM thaxtomin A, or 10 µM cellotriose. Error bars represent the standard error of the mean of twelve technical replicates (i.e. twelve separate plants in the same plate). **B** Plants were treated with 1 % methanol (MeOH), 1 % methanol + 10 µM thaxtomin A, or 10 µM cellotriose. Error bars represent the standard error of the mean of eight technical replicates.

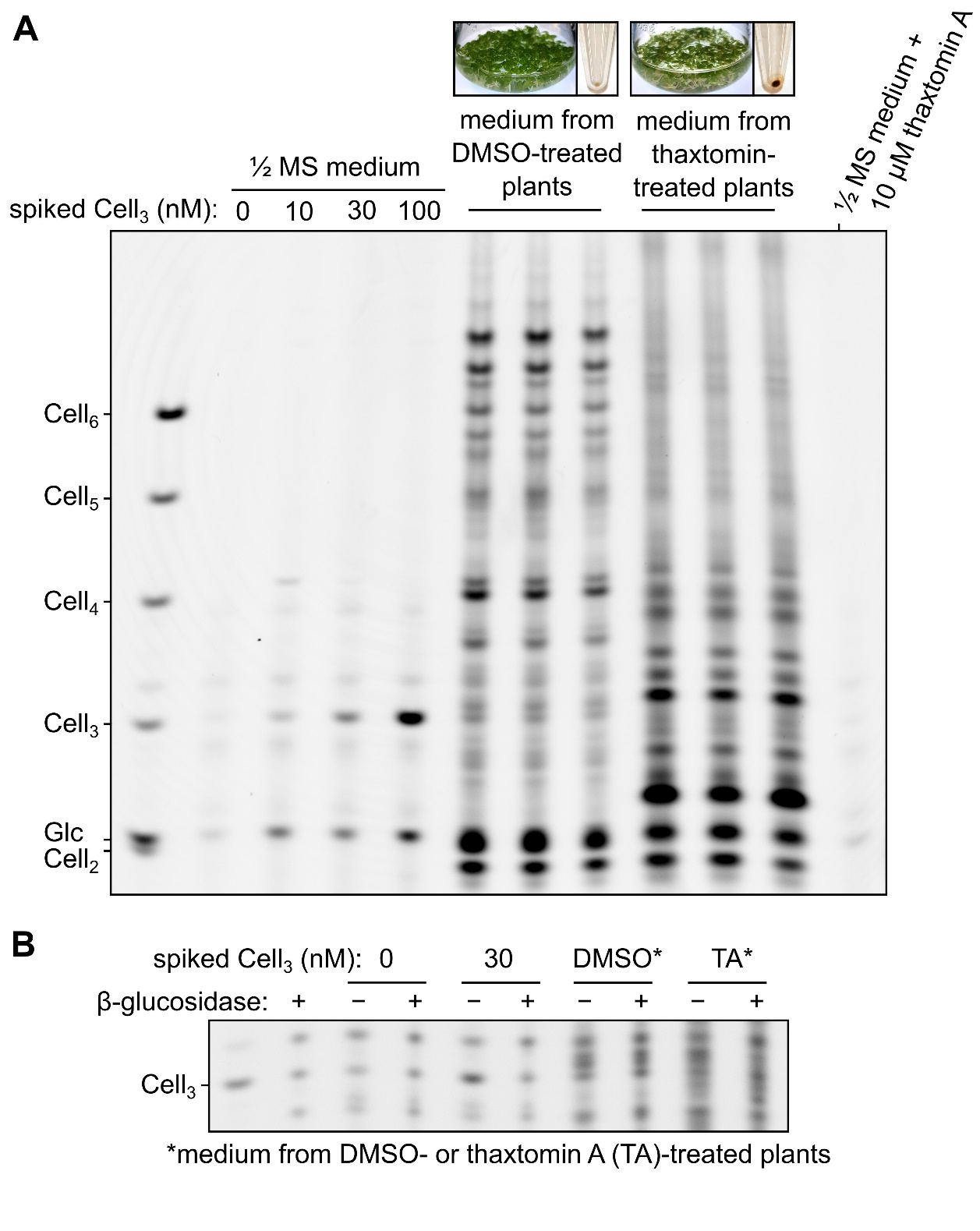

**Supplementary Fig. 16. Detection of oligosaccharides released from *N. benthamiana* seedlings grown in liquid medium**

**A** *N. benthamiana* seedlings were germinated and grown for nine days in liquid ½ MS medium. This medium was then replaced with fresh medium lacking sucrose but containing 0.5 % DMSO with or without 10 μM thaxtomin A. After three days, this medium was harvested and oligosaccharides were isolated by reverse phase chromatography. Control samples of medium spiked with cellotriose (Cell_3_) / thaxtomin A were subjected to the same isolation procedure. Oligosaccharides were then dried and covalently derivatised with 8-aminonaphthalene-1,3,6-trisulfonic acid (ANTS) fluorophore and subjected to polyacrylamide gel electrophoresis (polysaccharide analysis by carbohydrate electrophoresis; PACE). The appearance of seedlings (three separate flasks per treatment) after the three day treatment, as well as that of the dried eluate from chromatography, is shown above the gel. **B** Due to the presence of a background band co-migrating with cellotriose, the assignments of cello-oligosaccharide bands were verified by their sensitivity to β-1,4-glucosidase (lower gel). Purified oligosaccharides were digested with glucosidase prior to ANTS labelling and analysis by PACE. The difference in band intensity between untreated and digested samples represents the true abundance of cello-oligosaccharide.

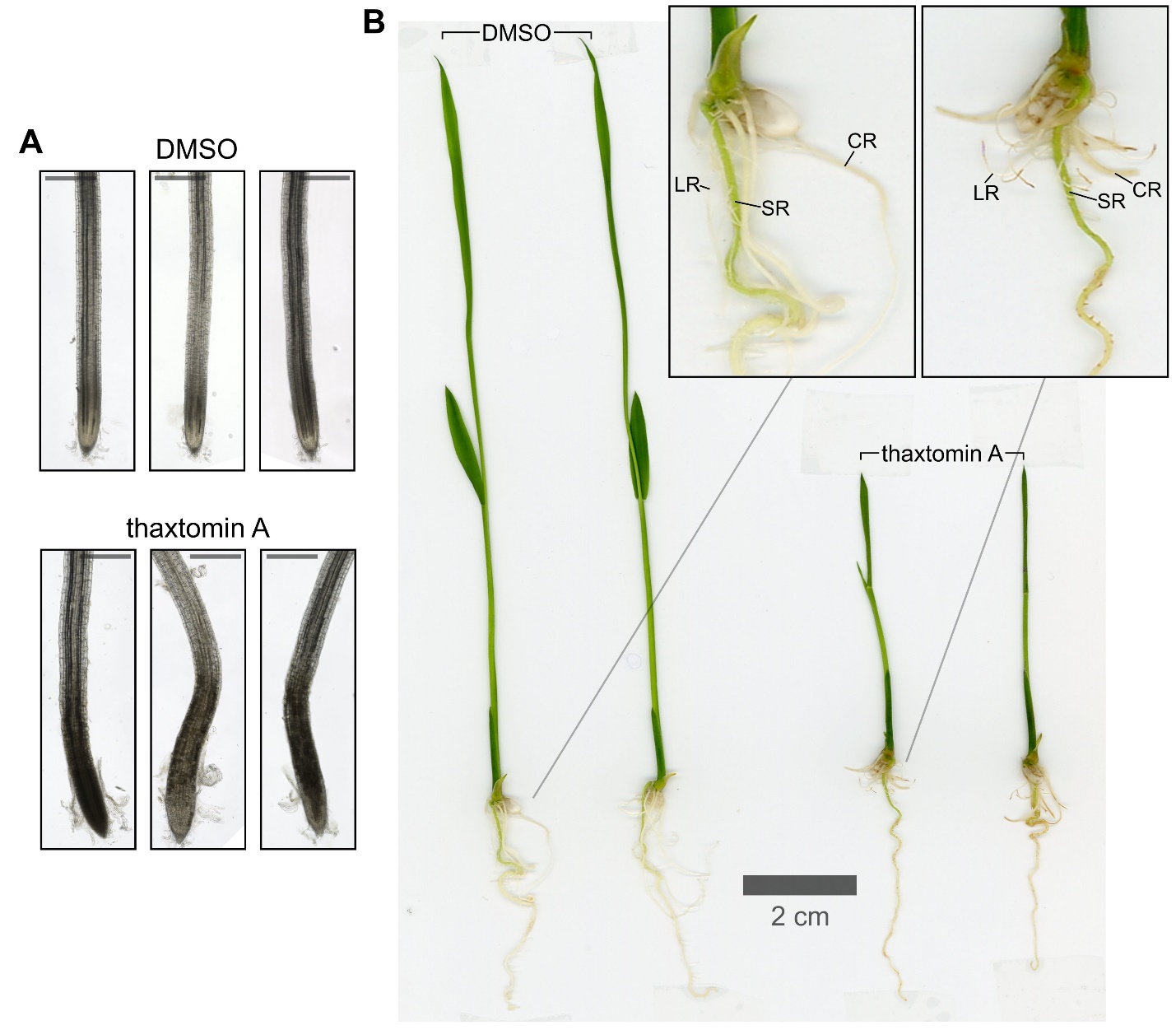

**Supplementary Fig. 17. Effects of thaxtomin A on rice seedling roots**

Rice seedlings were germinated on solid ½ MS. Five days after germination, plants were transplanted to liquid medium supplemented with 0.5 % DMSO ± 10 μM thaxtomin A. Five plants were analysed per treatment. **A** Appearance of lateral root tips 24 h after beginning of treatment. Scale bars: 300 μm. **B** Appearance of representative rice seedlings four days after beginning of treatment. Note discolouration of lateral roots (LR) and stunting of lateral roots and crown roots (CR). SR = seminal root.

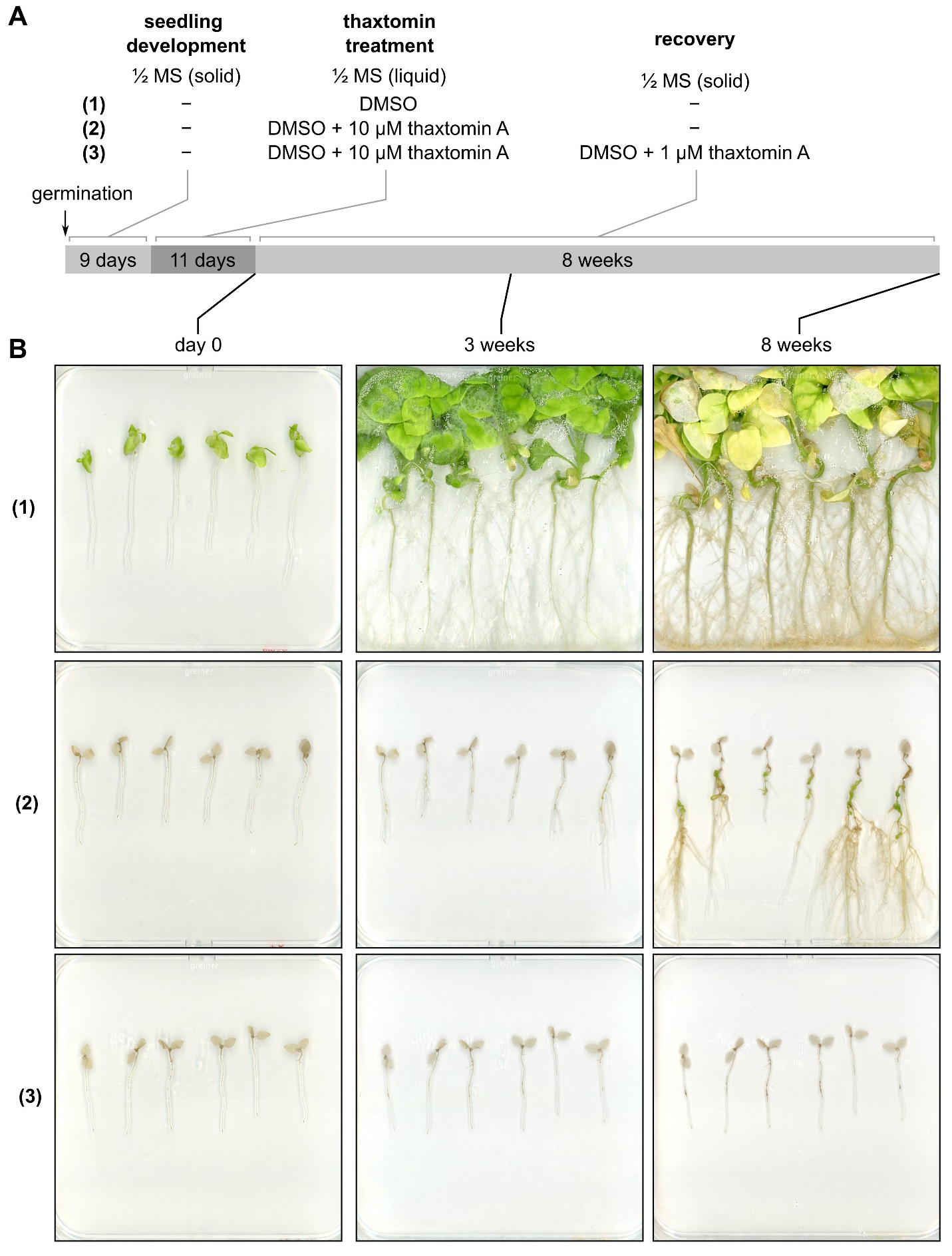

**Supplementary Fig. 18. Recovery of *N. benthamiana* seedlings after extended exposure to thaxtomin A.**

**A** Plants were treated according to the following scheme: Nine-day-old seedlings grown on solid ½ MS medium were submerged in liquid ½ MS medium containing 0.5 % DMSO lacking (1) or containing (2,3) 10 μM thaxtomin A and incubated for 11 days. The seedlings were then transplanted back to solid ½ MS medium, lacking (1,2) or containing (3) 0.5 % DMSO and 1 μM thaxtomin A. **B** The seedlings were then monitored for eight weeks. Close-ups of selected seedlings are shown in Figure 2.

**Supplementary Table 1. Cryo-EM data collection and processing parameters**

|  | ***Ptt*CesA8 +**  **thaxtomin A**  **PDB:** [**10DE**](https://www.rcsb.org/structure/10DE)  **EMD-**[**75083**](https://www.ebi.ac.uk/emdb/EMD-75083) | ***Ptt*CesA8 +**  **thaxtomin C**  **PDB:** [**10DF**](https://www.rcsb.org/structure/10DF) **EMD-**[**75084**](https://www.ebi.ac.uk/emdb/EMD-75084) | ***Ptt*CesA8 +**  **thaxtomin A**  **+ UDP-Glc**  **PDB:** [**10DG**](https://www.rcsb.org/structure/10DG) **EMD-**[**75085**](https://www.ebi.ac.uk/emdb/EMD-75085) | ***Ptt*CesA8 +**  **thaxtomin A**  **+ UDP**  **PDB:** [**10DH**](https://www.rcsb.org/structure/10DH) **EMD-**[**75086**](https://www.ebi.ac.uk/emdb/EMD-75086) | ***Ptt*CesA8**  **[H832W] +**  **thaxtomin A**  **PDB:** [**10DI**](https://www.rcsb.org/structure/10DI) **EMD-**[**75087**](https://www.ebi.ac.uk/emdb/EMD-75087) |
| --- | --- | --- | --- | --- | --- |
| **Data collection and processing** |  |  |  |  |  |
| Detector | Gatan K3 | Gatan K3 | Gatan K3 | Gatan K3 | Gatan K3 |
| Nominal magnification | 81,000 | 130,000 | 130,000 | 130,000 | 130,000 |
| Energy filter slit size (eV) | 10 | 10 | 10 | 10 | 10 |
| Voltage (kV) | 300 | 300 | 300 | 300 | 300 |
| Total electron dose (e^−^/Å^2^) | 50 | 60 | 60 | 60 | 60 |
| Spherical aberration (mm) | 2.7 | 2.7 | 2.7 | 2.7 | 2.7 |
| Target defocus range (μm) | −1.8–−0.8 | −1.6–−1.0 | −1.6–−1.0 | −1.6–−1.0 | −1.8–−1.0 |
| Calibrated pixel size (Å) | 1.065 | 0.652 | 0.652 | 0.652 | 0.652 |
| Movies collected | 6,991 | 8,900 | 8,538 | 11,108 | 10,746 |
| Symmetry imposed | symmetry expansion (C3) | symmetry expansion (C3) | symmetry expansion (C3) | symmetry expansion (C3) | symmetry expansion (C3) |
| Extraction box size (pixels) | 360 | 480 | 480 | 480 | 480 |
| Initial particle images (no.) | 3,225,422 | 1,241,662 | 913,889 | 1,938,387 | 2,121,236 |
| Final particle images (no.) | 455,288  (1,365,864 after expansion) | 120,830  (362,490 after expansion) | 188,949  (566,847 after expansion) | 207,709  (623,127 after expansion) | 257,586  (772,758 after expansion) |
| **Refinement** |  |  |  |  |  |
| Map resolution at FSC = 0.143 (Å) | 2.23 | 2.50 | 2.16 | 2.36 | 2.18 |
| Model composition |  |  |  |  |  |
| Non-hydrogen atoms | 16299 | 16287 | 16401 | 16518 | 16299 |
| Protein residues | 2016 | 2016 | 2022 | 2034 | 2016 |
| Non-protein atoms | 114 | 102 | 186 | 183 | 102 |
| *B* factors (mean, Å^2^) |  |  |  |  |  |
| Protein | 58.28 | 53.92 | 64.68 | 33.32 | 54.08 |
| Ligands | 49.06 | 39.25 | 58.93 | 17.78 | 42.31 |
| R.m.s. deviations |  |  |  |  |  |
| Bond lengths (Å) | 0.004 | 0.009 | 0.010 | 0.008 | 0.010 |
| Bond angles (°) | 1.081 | 1.216 | 1.231 | 1.194 | 1.231 |
| **Molprobity Statistics** |  |  |  |  |  |
| Clashscore | 1.05 | 1.05 | 0.95 | 1.37 | 1.08 |
| Ramachandran plot |  |  |  |  |  |
| Favored (%) | 97.88 | 97.12 | 97.13 | 97.46 | 97.42 |
| Allowed (%) | 2.12 | 2.88 | 2.87 | 2.54 | 2.58 |
| Outliers (%) | 0.00 | 0.00 | 0.00 | 0.00 | 0.00 |
| Rotamers |  |  |  |  |  |
| Favored (%) | 96.56 | 97.42 | 97.77 | 97.44 | 97.25 |
| Allowed (%) | 2.93 | 2.41 | 1.89 | 1.71 | 2.58 |
| Outliers (%) | 0.52 | 0.17 | 0.34 | 0.85 | 0.17 |
| Correlation coefficients |  |  |  |  |  |
| CC (mask) | 0.80 | 0.86 | 0.85 | 0.88 | 0.87 |
| CC (volume) | 0.79 | 0.83 | 0.84 | 0.84 | 0.85 |
| CC (peaks) | 0.60 | 0.63 | 0.61 | 0.72 | 0.66 |
| CC (box) | 0.60 | 0.62 | 0.61 | 0.70 | 0.65 |
